# Characterisation of *l(3)tb* as a novel tumour suppressor allele of *DCP2* in *Drosophila melanogaster*

**DOI:** 10.1101/574707

**Authors:** Rakesh Mishra, Rohit Kunar, Lolitika Mandal, Debasmita P Alone, Shanti Chandrasekharan, Anand Krishna Tiwari, Ashim Mukherjee, Madhu G Tapadia, Jagat Kumar Roy

## Abstract

Mutants provide an excellent platform for the discovery and characterization of gene functions. The present communication is a pioneering treatise on a hitherto undescribed function of the gene coding for the mRNA decapping protein 2 (*DCP2*) in *Drosophila melanogaster. DCP2*, the gene coding for the mRNA decapping enzyme, has been studied in various model organisms in the light of maintenance of transcript abundance and stability but has never been implicated in tumourigenesis. Herein, we describe the mapping and characterization of a novel tumour suppressor allele of *DCP2* (CG6169), which we named as *lethal(3)tumorous brain* [*l(3)tb*]. The homozygous mutant individuals show prolonged larval life, develop larval brain tumors and are lethal in the larval/pupal stages. The tumour is characterized by the presence of increased number of superficial neuroblasts, abnormal chromosomal condensation and causes overgrowth in the wing and the eye-antennal discs of the homozygous mutant larvae, all of which are rescued by the introduction of a functional copy of *DCP2* in the mutant background, thereby establishing the causal role of the mutation and providing a genetic validation of the allelism. Our findings therefore ascribe a novel role of tumor suppression to *DCP2* besides its cognate function of mRNA decapping and thereby identify it as a potential candidate for future research on tumorigenesis.

## Introduction

Tumorigenesis occurs either due to gain-of-function of an oncogene or loss-of-function of a tumor suppressor gene. As many as 57 oncogenes and 81 tumor suppressor genes have been identified through genome wide sequence studies apart from conventional approaches of various intragenic mutations (Vogelstien *et al*. 2013). Collective studies from developmental biology in the field of *Drosophila*, mouse and humans revealed that in most cases, the initiating event in the formation of a malignant tumor or neoplasia is a loss of function in the regulatory genes controlling cell growth and differentiation (Gateff and Schneiderman 1967, 1969; Harris *et al*. 1969; Knudson *et al*. 1971; Harris 2005; Papagiannouli and Mechler 2013). *Drosophila* shows classic hallmarks of cancer, such as evasion of apoptosis, sustained proliferation, metastasis, prolonged survival, genome instability and metabolic reprogramming (Luo *et al*. 2009; Hanahan and Weinberg, 2011) and these phenotypes result from loss of function of a tumor suppressor gene in general, also called as recessive oncogenes (Mechler 1994).

In our present communication, we show the characterization of a new tumor suppressor mutation in *Drosophila melanogaster*, which was subsequently identified to be an allele of *Decapping protein 2* (*DCP2*; CG6169). The homozygous mutant individuals show prolonged larval life, develop larval brain tumors and are lethal in the larval/pupal stages. Hence, the mutation was initially named as *lethal(3)tumorous brain* [*l(3)tb*]. The tumor is characterized by the presence of increased number of superficial neuroblasts and abnormal chromosomal condensation. The mutation is also responsible for causing overgrowth in the wing and the eye-antennal discs of the homozygous mutant larvae. Recombination and complementation mapping of the mutation show that it is allelic to *Decapping protein 2* (*DCP2*) located at 72A1 on the left arm of chromosome 3. Complementation analyses of the mutation with alleles of *DCP2* show phenotypes similar to *l(3)tb* homozygotes, thereby confirming the proposed allelism. Over-expression of wild type *DCP2* in the mutant background rescues the mutant phenotypes, thereby providing a genetic validation of allelism. Molecular mapping identified the mutation to be residing in the 5’UTR coding region of *DCP2*. Our findings therefore ascribe a novel role of tumor suppression to *DCP2* besides its cognate function of mRNA decapping and thereby identify it as a potential candidate for future research on tumorigenesis.

## Materials and Methods

### Fly strains and rearing conditions

Fly cultures were raised on standard food containing agar, maize powder, yeast, sugar and supplemented with anti-fungal (Nepagin, methyl-p-hydroxy benzoate) and anti-bacterial (propanoic acid) chemicals at 23+1°C. Subsequently genetic crosses were carried out following standard procedures. Oregon R+ was used as the wild type strain. The recessive *l(3)tb* mutation (*yw*; +/+; *l(3)tb /TM6B, Tb*^1^, *Hu, e*^1^) was isolated in a genetic screen and the mutation was maintained with the *TM6B* balancer which established its linkage to chromosome 3. The multiply marked “*rucuca*” (*ru h th st cu sr e ca/TM6B, Tb*) and “*ruPrica*” (*ru h th st cu sr e Pr ca/TM6B,Tb*) chromosomes were employed for recombination mapping (Lindsley and Zimm 1992). *w; Δ2 −3, Sb/TM6B,Tb*^1^, *Hu, e*^1^ (Cooley *et al*. 1988) and *CyO, P{Tub-Pbac/T}2/Wg[Sp-1];+/TM6B, Tb, Hu,e^1^* (Bloomington Stock No. 8586) were used for providing transposase source for *P* element and *piggyBac* specific transposable element, respectively, in mutagenesis experiment. Second chromosome balancer *Sp/CyO* (O’Donnell *et al*. 1975) and third chromosome balancer yw; *TM3, Sb, e/ TM6B, Tb^1^, Hu, e^1^* were obtained from Bloomington *Drosophila* stock center. The *elav-GAL4* (Lin and Goodman 1994), *y^1^ w; P{Act5C-GAL4}25F01/CyO, y w; +/+; Tub-GAL4/ TM3, Sb, e* and *UAS-GFP* stocks were obtained from the Bloomington *Drosophila* Stock Center. Transgenic *UAS-DCP2 RNAi* (on chromosome 1 and chromosome 2 were obtained from Vienna *Drosophila* RNAi stock Centre, VDRC and *w; UAS-mCD8::GFP* (Lee and Luo, 1999) from Bloomington *Drosophila* stock center. The lethal insertion mutants of gene *Decapping protein2 - PBac{RB}DCP2^e00034^/TM6B,Tb^1^Hu, e^1^* (Thibault *et al*. 2004) and *P{GT1}DCP2^BG01766^/TM3, Sb^1^, e^1^* (Lukacsovich *et al*. 2001) were obtained from Exelixis Stock Center, Harvard University and Bloomington *Drosophila* stock center, respectively.

Deficiency stock *Df(3L)RM96* was generated in the laboratory using progenitor *P* element stocks, viz., *P{RS5}5-SZ-3486, P{RS5}5-SZ-3070, P{RS3}UM-8356-3, P{RS3}UM-8241-3, P{RS3}CB-0072-3, yw P{70FLP,ry^+^}3Fiso/y^+^Y;2iso;TM2/TM6C,Sb, w^1118^iso/y^+^Y;2iso;TM2/TM6C,Sb*, obtained from Vienna *Drosophila* Resource Center (Golic and Golic, 1996; Ryder *et al*. 2007). Various deficiency stocks (Table S1, S2) and transposon insertion fly stocks (Table S3) used for complementation analysis were obtained from Bloomington *Drosophila* stock centre and Exelixis stock center.

### Analysis of lethal phase in *l(3)tb* homozygotes

For analysis of lethal phase and morphological anomalies associated with the homozygous *l(3)tb* mutation, embryos were collected at the intervals of 2h on food filled Petri-plates. Embryos from wild type flies were collected as controls. The total number of eggs in each plate was counted and the embryos were allowed to grow at 23°C or 18°C or 16°C (+1°C). Hatching of embryos and further development of larval stages was monitored to determine any developmental delay. Mutant larvae, at different stages, were dissected and the morphology of larval structures was examined.

### Meiotic recombination mapping of *l(3)tb* mutation

Genetic recombination with multiple recessive chromosome marker *ru cu ca* was performed to map mutation in *yw*: +/+; *l(3)tb/TM6B, Tb* mutant in order to map the mutation. The *y w; l(3)tb/TM6B* males were crossed to virgin +/+; *ru Pri ca/TM6B* females to recover *l(3)tb* without *y w* on X-chromosome. The F1 *l(3)tb/TM6B* males were crossed to virgin +/+; *ru cu ca* females and the F2 progeny +/+; *l(3)tb/rucuca* virgin females were selected. Recombination of the *l(3)tb* and *rucuca* bearing chromosomes is expected to occur in these F2 flies. These F2 virgins were then crossed to *ru Pri ca/TM6B* males to score the frequency of recombinants in the F3 progeny. The *l(3)tb* phenotype cannot be scored in these flies because in F3, these flies carry *l(3)tb* mutation only in the heterozygous condition. Therefore, all the F3 progeny males obtained, were individually scored for *ru, h, th, st, cu, sr, e* and *ca* phenotypes and then they were individually crossed with 2-3 virgin *l(3)tb/TM6B* females to identify which of them had the *l(3)tb* mutation along with other scored markers.

### Complementation mapping of the *l(3)tb* mutation

Complementation analysis of the mutation in *l(3)tb* allele was carried out with Exelixis and DrosDel deficiency stocks spanning the entire chromosome 3 (Table S1). Virgin *y w*; +/+; *l(3)tb/TM6B, Tb* females were crossed with the males of the various deficiency stocks and the non-tubby F1males heterozygous for *l(3)tb* and the deficiency were observed carefully for the lethal phenotype. The lethal molecular lesion in *l(3)tb* was also checked for their allelic partners directly by complementation analysis using molecularly characterized lethal *P*-insertion alleles. 25 lethal *P*-insertion alleles were selected from the region narrowed down through recombination and deficiency mapping. Genetic crosses were set taking males from the lethal *P*-insertion and virgin females from the mutant *l(3)tb* fly stock. The non-tubby progeny, heterozygous for lethal P-insertion and lethal *l(3)tb* mutation were scored for the phenotype.

Reversion analysis was performed by the excision of *piggyBac* transposon in *DCP2^e00034^* with the help of *piggyBac* specific transposase source, *CyO, P{Tub-Pbac}2/Wg_SP-1_* (Thibault *et al*. 2004) and similarly by the excision of *P*-element in *DCP2^BG01766^* strain using Δ*2-3,Sb/TM6B, Tb^1^, Hu, e^1^* (Cooley *et al*. 1988) transposase source as ‘jumpstarter stock’, both obtained from Bloomington stock centre. Virgin flies from the mutator stocks *DCP2^e00034^* and *DCP2^BG01766^* strain were crossed to male flies from respective ‘jumpstarter stock’. F1 male flies with mosaic eye pigmentation carrying both the transposase and respective transposons were selected and crossed to JSK-3 (*TM3, Sb, e^1^/TM6B, Tb^1^, Hu, e^1^*) virgins and from the next generation rare white eyed revertant F2 flies were selected (Figure S9).

### Genomic DNA extraction, Primer Design and PCR

Single Fly genomic DNA isolation for PCR was done as essentially described (Gloor and Engels 1992; Garozzo and Christensen 1994). Genomic DNA for polymerase chain reaction (PCR) was isolated by homogenizing 50 male flies from each of the desired genotype or 80-100 third instar larvae from homozygous mutant *l(3)tb* (Sambrook *et al*. 1989).

Primers were designed by using a modified version of the Primer3Plus. Sequence for the genomic region, narrowed down by the mapping strategies, was downloaded from the FlyBase (version R5). To analyze any molecular lesion in the DNA of homozygous *l(3)tb* mutant, 28 pairs of primers were designed for genomic region of gene *Decapping protein 2* (CG6169; 7.69 kbp) from 3L:15811834.15819523 (Table S5, S6, S7, S8) (Rozen and Skaletsky 2000).

Genomic DNA was subjected to PCR, with final volume of 25 μl containing 25 pM each of the two primers, forward and reverse, for each primer pair, 200 μM of each dNTP (New England Biolabs, USA) and 1.5U of *Taq* DNA Polymerase (New England Biolabs, USA). The cycling parameters included an initial denaturation of 5 min at 95°C followed by 30 cycles of denaturation at 94°C for 1 min, annealing (temperature accordingly to the specific primers for 1 min) and extension at 72°C for 1 min, with 10 min extension in the last cycle. PCR products were checked for amplification using 2% agarose gel along with 100bp DNA ladder or the *pUC12* vector DNA digested with *HinfI* as a molecular marker.

### Automated Sequencing of PCR amplicons

The PCR products were sequenced directly with the help of Applied Biosystem 3130 Genetic Analyser platform, BBI, USA. The PCR products were eluted from 0.8% agarose gel using the gel-extraction kit (Fermentas Life Sciences, EU), following manufacturer’s protocol and were processed by Applied Biosystems cycle sequencing kit version 3.1 using ready reaction (RR) mix and 10X PCR buffer in a 10 μl reaction volume and processed in both forward and reverse directions with the same set of primers employed for initial PCR. The fluorescently labeled DNA product was precipitated using ABI Big Dye Terminator Clean up method following manufacturer’s recommendation and dissolved in Hi-Di (Formamide) and processed for sequencing. The sequences were base-called and assembled using Geospiza FinchTV version 1.4 (http://www.geospiza.com/Products/finchtv.shtml). Homology search and alignments were performed using BLAST algorithm available at NCBI and FlyBase.

### RNA isolation and Reverse Transcription-PCR

Total RNA was isolated from healthy wandering third instar larvae of wild type and delayed third instar larvae from homozygous *l(3)tb* mutant using TRIzol reagent following the manufacturer’s recommended protocol (Sigma-Aldrich, India). The samples were incubated with 2U of RNase-free DNaseI (MBI fermentas, USA) for 30 minutes at 37°C to remove any residual DNA and dissolved in DEPC (Diethyl pyrocarbonate, Sigma, USA) treated water. First strand cDNA was synthesized using M-MuLV Reverse Transcriptase (RT) (Life Technologies, Invitrogen). Briefly, ~5 μg of total RNA using 20U of RNase OUT (Invitrogen), 80 pmol of oligo-dT_18_ primer (New England Biolabs, USA), 500μM of dNTP mix and 100U of M-MuLV reverse transcriptase (SuperScript III Reverse Transcriptase, Invitrogen) were added to a final reaction volume of 20μl followed by incubation for 1h at 37°C. The reverse transcriptase enzyme was inactivated at 65°C for 15 min. 1/20^th^ (1μl) volume of the reaction mixture was subjected to second strand synthesis using the primer pair for the gene *DCP2* (3L: 15814923-15815518) giving an amplicon size of 595 bp with genomic DNA (gDNA) and 539 bp with complementary DNA (cDNA). G3PDH (Glycerol 3 Phosphate dehydogenase) was used as internal control. The specific primers used were: Forward (DCP2): TATCAAATCCATGCCCGTTG and Reverse (DCP2): GTCACAGGAGTGCGAAATGA. Forward (G3PDH): 5’-CCACTGCCGAGGAGGTCAACTA-3’; Reverse (G3PDH): 5’-GCTCAGGGTGATTGCGTATGCA-3’. The thermal cycling parameters included an initial denaturation at 94° C (4 min) followed by 30 cycles of 35 sec at 94°C, 30 sec at 60°C (For G3PDH), 40 sec at 54°C (for DCP2), 1 min at 72 °C. Final extension was carried out at 72°C for 5 min. The PCR products were electrophoresed on a 2% agarose gel with appropriate molecular weight markers.

### Poly-acrylamide gel electrophoresis (PAGE) and Immunoblotting

The larval brain ganglia of the desired genotypes were dissected out in PSS and homogenized in the protein sample buffer (100 mM Tris, pH 6.8; 1M DTT; 10% SDS; 100 mM phenylmethyl sulphonyl fluoride (PMSF), pH 6.8; 1% bromophenol blue and 1% glycerol). Protein samples were resolved in denaturing condition in 12% vertical SDS-polyacrylamide slab gel using a discontinuous buffer system (Laemmli 1970) and electrophoretically transferred to polyvinylidene fluoride membrane (PVDF, Millipore, USA) at 0.8mA/cm^2^ or 50V through wet blotting apparatus (Biotech, India). The membrane was rinsed 2 times for 5 min each, in TBST (100 mM Tris, pH 7.4; 150 mM NaCl, 0.1% Tween 20) and blocked for 2 h at RT in blocking buffer (5% skimmed milk powder in TBST). The membrane was probed with primary antibody against Dlg (1:100) and detected with Anti mouse-HRP secondary antibody (dilution 1:2000) using enhanced chemiluminescence (ECL) detection as per manufacturer’s instruction (SuperSignal, Pierce, USA). Membrane was incubated in 100 mM β-mercapto ethanol, 2% SDS, 62.5 mM Tris, pH 6.8 at 50°C for 30 min and probed with anti β-tubulin antibody at 1:200 dilutions.

### Cytological techniques

Mitotic chromosomes from brain ganglia of *l(3)tb* homozygous larvae of different ages (5^th^ to 12^th^ day after hatching) and from that of wild type late third instar larvae were prepared as per standard protocol (Lakhotia *et al*, 1979). Brain ganglia from mutant larvae of different ages and wild type late third instar larvae were dissected in Poel’s salt solution, incubated in 1% toludine blue (pH 7) for 1h at 24°C, fixed in Bodain’s fixative and then processed as mentioned by Truman and Bate (1988) to visualize and score the number of darkly stained superficial neuroblasts.

### Analysis of Eye morphology

The fly was anaesthetized, and decapitated with a sharp blade or needle. The decapitated head was briefly dipped in a drop of transparent nail polish. The head was then placed on a clean, dry area of the same slide and the nail polish layer was allowed to dry at RT for 5-10 min. The dried layer of nail polish was peeled off from the eye with the help of fine dissecting needles and was carefully placed on another clean glass slide with the imprint side facing up and flattened by gently placing a cover slip. The eye imprint was then examined under a microscope using 20X differential interference contrast (DIC) objective as described (Arya and Lakhotia 2006).

### BrdU incorporation studies to study replication profile of neuroblasts

Brain ganglia from larvae of different age groups, i.e., 6 days ALH in case of wild type and 6 to 13 days ALH in case of *l(3)tb* homozygous larvae were dissected in Poels’ salt solution, labeled with Bromodeoxy-Uridine (BrdU) Sigma, 20μM) at room temperature for 60 min in dark. They were then fixed in 90% ethanol for 30 min and passed through descending ethanol grades (15 min each), briefly hydrolyzed in 2N HCl for 15 min at room temperature, washed in PBS and incubated with 10% normal goat serum for 1 h at 4°C for blocking and incubated overnight with anti-BrdU antibody at 4°C. The bound anti-BrdU antibody was subsequently detected by anti-mouse IgG-FITC conjugate (Sigma, dilution 1:64) under a fluorescence microscope after mounting the ganglia in antifade (Sigma).

### Immunostaining

The imaginal discs and/or brain ganglia were collected from wild type *Oregon R*^+^ wandering 3^rd^ instar larvae, just before pupation (110 h, AEL) and in mutant homozygous *l(3)tb* from day 6 and day 10/12, as homozygous mutant larvae has delayed development up to 12-13 days. The tissues were processed for immunostaining with desired antibodies as described (Banerjee and Roy, 2017). For imaging endogenous GFP expression, tissues were dissected in 1X PBS and rinsed with 0.1% PBT. Counterstaining was performed with either DAPI (4’, 6-diamidino-2-phenylindole dihydrochloride, Sigma) at 1μg/ml, or phalloidin-TRITC (Sigma-Aldrich, India) at 1:200 dilutions. Tissues were mounted in DABCO (antifade agent, Sigma). Samples were examined with Zeiss LSM 510 Meta laser scanning confocal microscope at appropriate settings using plan apo 20X, 40X or 63 X oil immersion objectives.

### Antibodies

Primary antibodies used in this study were - Anti-Discs large, 4F3 (Dilution 1:50, Developmental Studies Hybridoma Bank, Iowa, USA), Anti-Armadillo (1:100, a kind gift by Prof LS Shashidhara, Pune, INDIA), Anti-Elav (Rat-Elav-7E8A10, Dilution 1:100, DSHB, USA), 22C10 (Dilution 1:100, DSHB, Iowa, USA), Anti-DE-Cadherin (DCAD2,Dilution 1:20, DSHB, Iowa, USA), Anti-phospho-Histone 3 (Dilution1:500, Millipore, Upstate, USA), Anti-Fasciclin II (1D4, Dilution 1:50, DSHB, Iowa, USA), Anti-Deadpan (Dilution 1:800, a kind gift from Prof. Volker Hartenstein, University of California, USA), Anti-repo (8D12, Dilution 1:10, DSHB, Iowa, USA), and Anti-β-tubulin (E7, Dilution 1;200, DSHB, Iowa, USA). Secondary antibodies used were Alexa Fluor 488 conjugated goat anti-mouse IgG, Alexa Fluor 488 conjugated goat anti-rat IgG, Alexa Fluor 488 conjugated donkey anti-rabbit IgG and Alexa Fluor 488 conjugated goat anti-guinea pig IgG from Molecular Probes, USA at a dilution of 1:200. Cy3 conjugated Anti-rabbit IgG and Cy3 conjugated Anti-mouse IgG (Sigma-Aldrich, India) were also used at 1:200 dilutions, biotinylated anti-rabbit IgG (Vector Lab) and streptavidin conjugated HRP (Vector Lab) and Anti-mouse HRP (Bangalore Genie, India). The immunostained slides were observed under Zeiss LSM 510 Meta Laser Scanning Confocal microscope, analysed with LSM softwares and assembled using Adobe Photoshop 7.0

### Statistical analysis

Sigma Plot (version 11.0) software was used for statistical analyses. All percentage data were subjected to arcsine square-root transformation. For comparison between the control and experimental samples, One-Way ANOVA was performed. Data were expressed as mean ± S.E. of mean (SEM) of several replicates.

## RESULTS

### *l(3)tb* homozygotes show the classic hallmarks of cancer in *Drosophila* including developmental delay, abnormal karyotype, larval/pupal lethality alongwith tumorous brain and wing imaginal disc

Developmental analysis of *l(3)tb* homozygotes showed that while embryos hatched normally and developed alike their heterozygous siblings [*l(3)tb/TM6B*], the third instar larvae reached the wandering stage quite late with the larval stage extending up to 12 or 13 days (**Figure 1B**). Only 66.8% of the larvae survived to pupate (**Table 1**), but died in the pupal stage following bloating, enhancement in size and cessation of growth (**Figure 1A**). Hence, the mutation is absolutely lethal with the lethality being pronounced in the pupal stage. Lowering the temperature to 16°C or 18°C reduced the larval mortality, causing 96% of larvae to pupate but did not improve pupal survival (**Figure 1C and D**). Analysis of larval brain and imaginal discs in the homozygotes in the early (Day 6) and late (Day 10-12) larval phase showed gross morphological alterations in the size of the larval brain, wing and eye imaginal discs (**Figure 2A–G**) as compared to the wild type counterparts of similar developmental stage (115h ALH; **<u>A</u>**fter **<u>L</u>**arval **<u>H</u>**atching). The brain was smaller in size than the wild type (*Oregon R*^+^) or heterozygous [*l(3)tb/TM6B*] individuals till 115 ALH but started showing aberrant growth in the dorsal lobes thereafter, showing significant differences in the diameter and area of the lobes. The overgrown brain hemispheres remained more or less symmetric in most of the cases, except in some where it got deformed and fused with the imaginal discs (**Figure 2J and K**). A similar trend in morphological aberration was observed in the wing discs, which remained smaller initially but enlarged sufficiently later (**Figure 2L**), with abnormal protrusion in the wing pouch.

**FIGURE 1.**
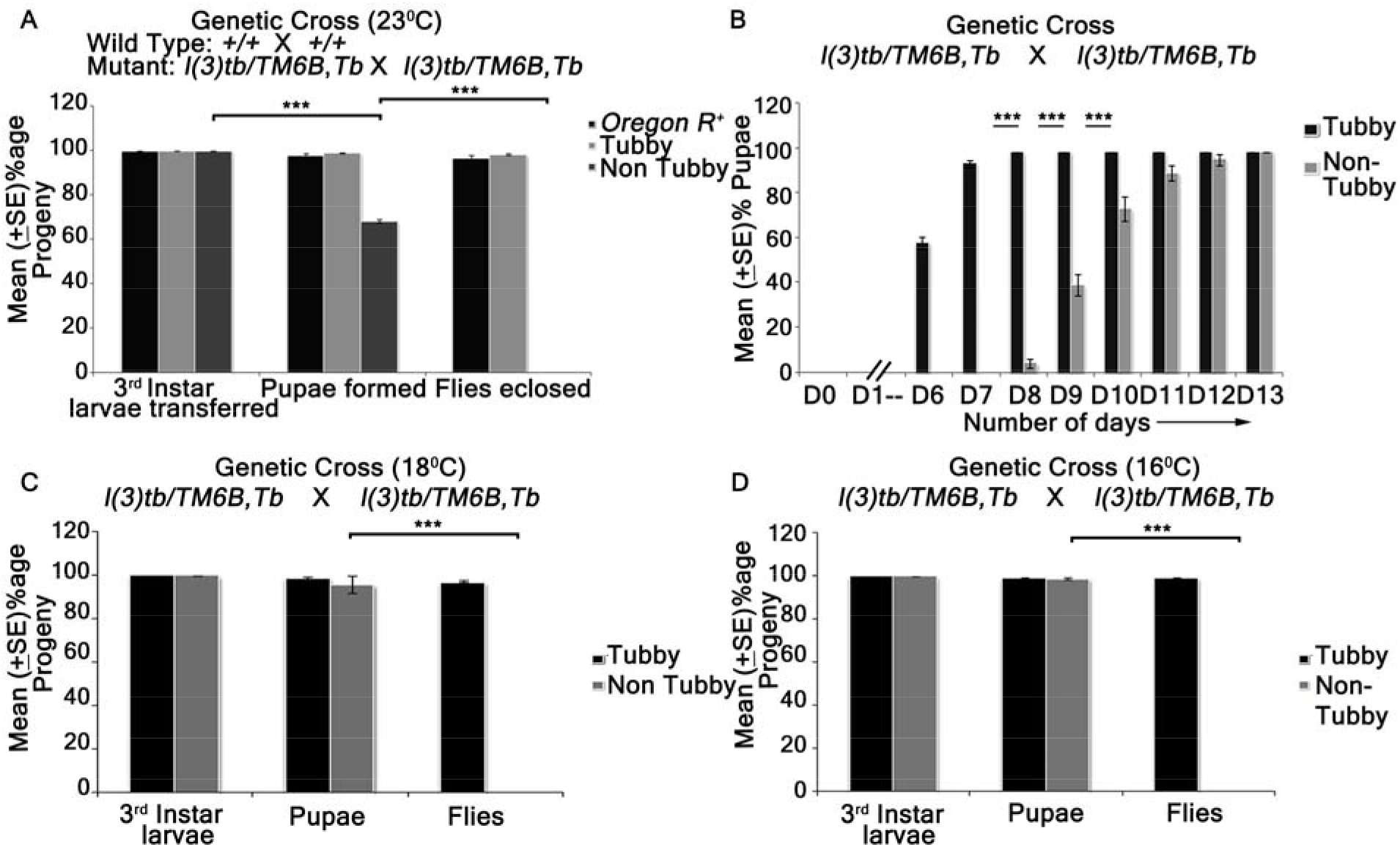
Homozygous *l(3)tb* show delayed larval development with lethality at larval/pupal stage (A, B) and is not a conditional temperature sensitive allele (A, B, C). Homozygous *l(3)tb* progeny, at 23°C, showed lethality at larval and pupal stages and no flies eclosed as compared to wild type and heterozygous *l(3)tb* progeny with balancer chromosome (A). Homozygous *l(3)tb* progeny individuals demonstrated extended larval life up to day 12/13 where as heterozygous progeny individuals followed the normal wild type pattern of development (B). (C) and (D) show significant increase in viability of homozygous (non-tubby) *l(3)tb* larvae at lowered temperatures of 18°C and 16°C respectively, though there also occurred absolute lethality at pupal stages. Each bar represents mean (±S.E.) of three replicates of 100 larvae in each. *** indicates p<0.005 *** indicates p<0.005.

**TABLE 1.**
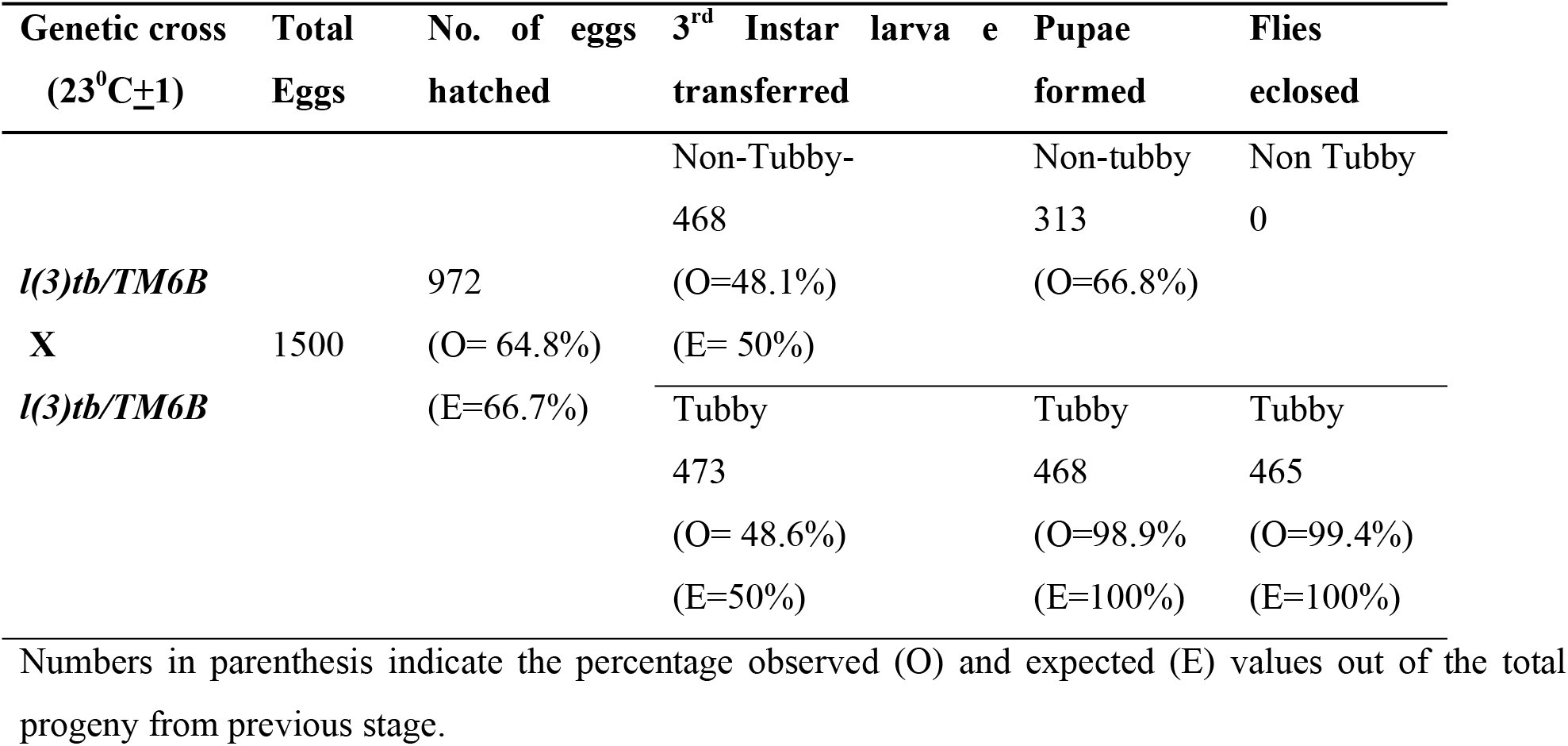
Homozygous mutation in *l(3)tb* causes larval and pupal lethality.

Analysis of mitotically active cell population by screening for the metaphase marker protein, phosphorylated histone H3 (PH3) revealed increased number of active mitoses in the mutant homozygous brains (**Figure 3A-O; 3V**) and wing discs (**Figure 3P-V**) (Day 6) in comparison to the wild type, the number of which increased with increase in larval age of the mutant. However, mitotic karyotypes of the mutant brain lacked numerical aberrations, despite showing extensive variability in condensation (**Figure 2H and I**).

**FIGURE 2.**
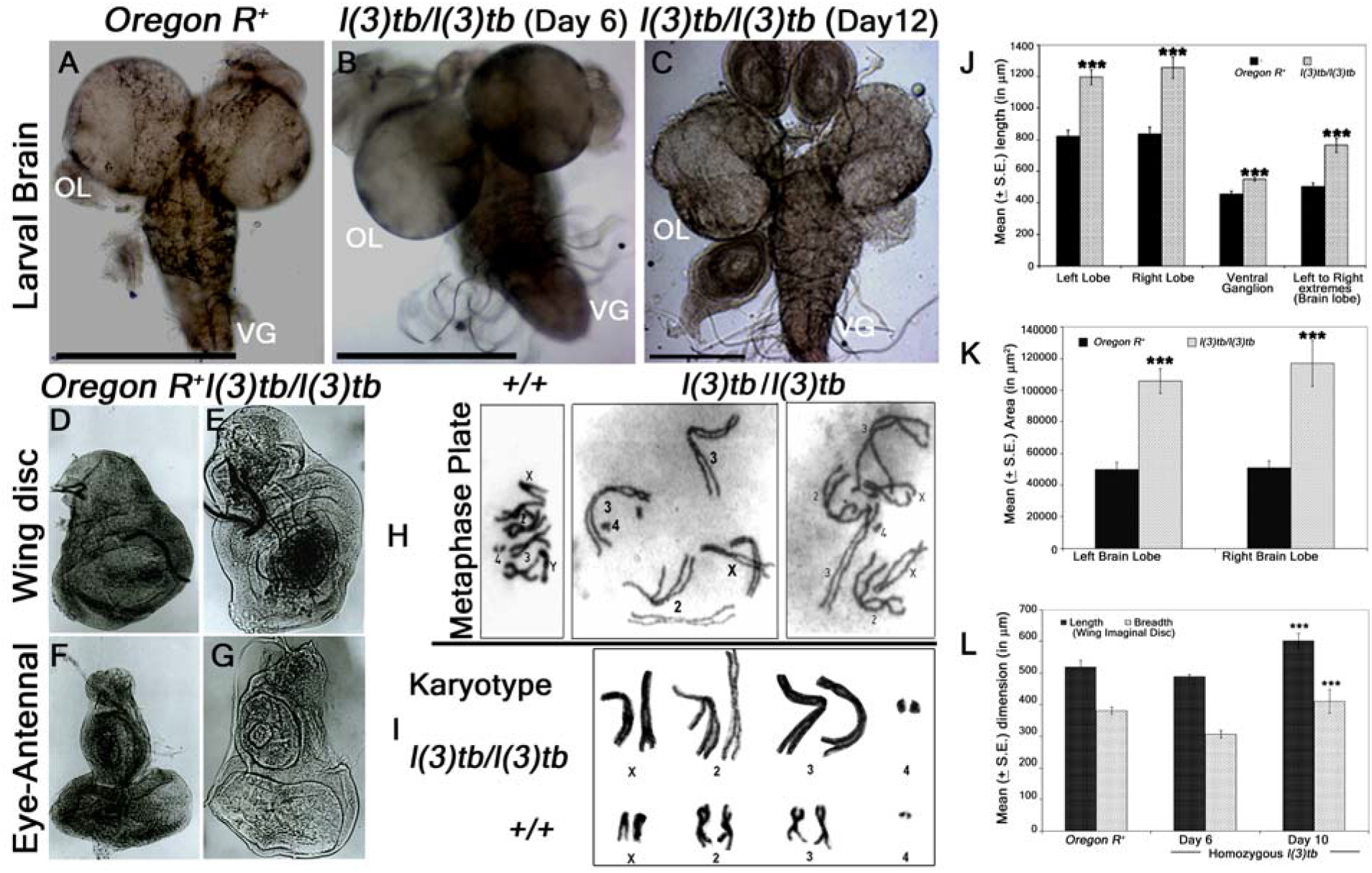
Homozygous *l(3)tb* mutants show severe morphological alteration in delayed 3^rd^ instar larval brain, wing and eye-antennal disc of. Homozygous *l(3)tb* mutant 3^rd^ instar larvae revealed tumorous brain of day 12 (C) as compared to day 6 of homozygous mutant (B) and day 5 of wild type, *Oregon R*^+^ (A). *l(3)tb* homozygotes exhibited highly significant differences in the overall circumference of the left and right brain lobes in the delayed stage (day 10) as compared to the respective wild type brain lobes (J). Significant differences were found in the area (μm^2^) of respective brain lobes of *l(3)tb* homozygotes and wild type (K). Dimension of wing and eye-antennal imaginal discs of delayed 3^rd^ instar larvae from homozygous *l(3)tb* mutant revealed significant increase in size (D,E,F,G). Length and breadth of wing discs from 3^rd^ instar larvae of *l(3)tb* mutant of day 6, was found to be smaller than the wing imaginal discs from wild type, but wing discs from extended larval period (day 10) showed significant increase in the size (L). Metaphase chromosome preparation of brain cells (H) from wild type and *l(3)tb* homozygotes exhibited abnormal karyotypes (I) where *l(3)tb* homozygotes showed less condensed and extended chromosome morphology as compared to wild type, *Oregon R*^+^. *** denotes p<0.005

**FIGURE 3.**
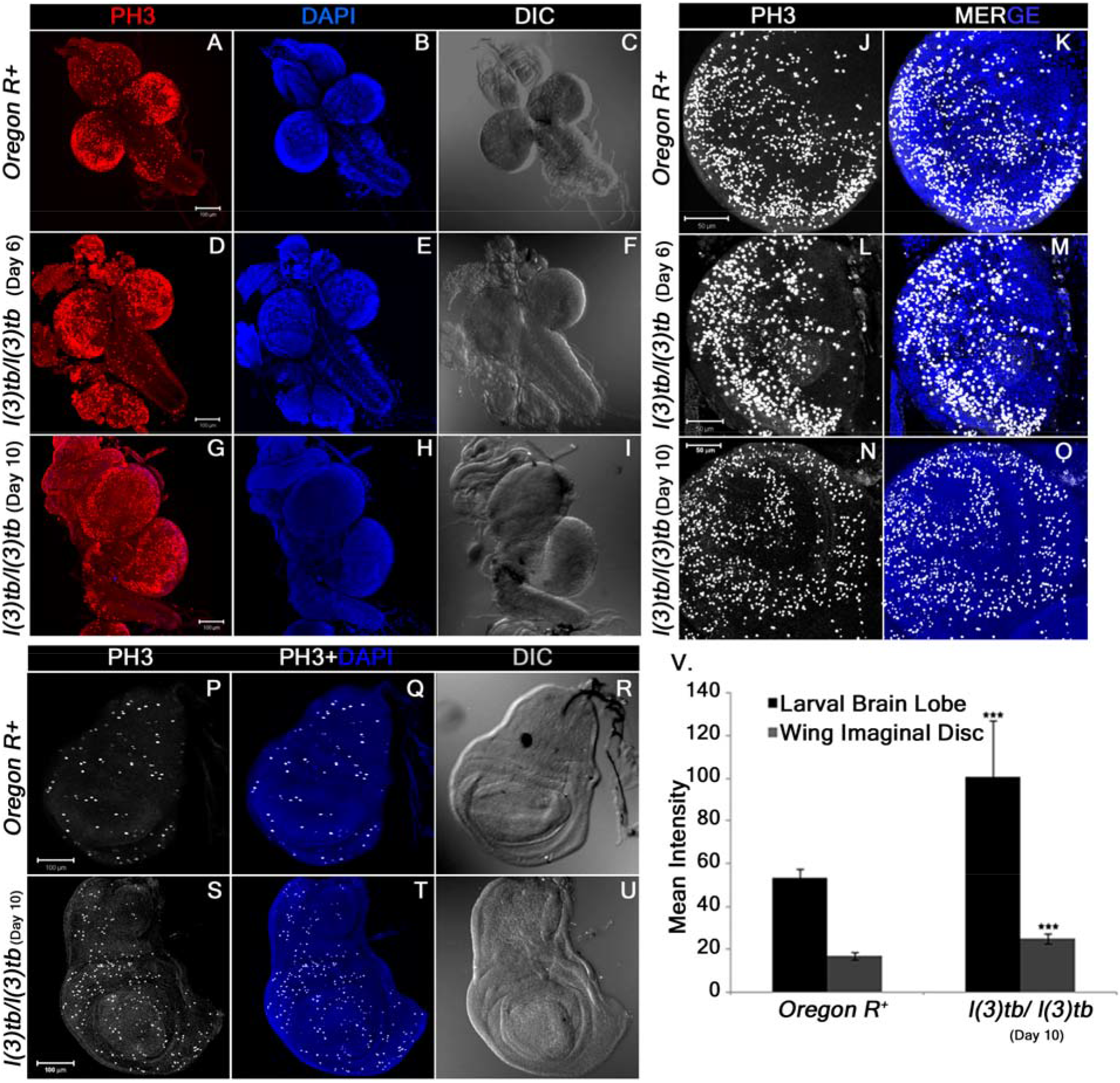
Enhanced mitotic potential observed in the tumourous tissues of homozygous *l(3)tb* as shown in larval whole brain (A), brain lobes (D, G) and wing imaginal discs (S) immunostained with phosphor-histone 3 (PH3), a potent mitotic marker. Distribution of PH3 labeled cells counter stained with DAPI cells in wild type (A) and homozygous *l(3)tb* (Day 6 and Day 10) larval brain (D, G) and also in wild type brain lobes (B, C) and homozygous mutant brain lobes (E, F for day 6; H, I for day 10) indicated high mitotic index as compared to wild type. Similarly, more mitotic positive cells were seen in tumorous wing imaginal discs (day 10) of homozygous mutant l(3)tb (II-D,E) as compared to wild type, *Oregon R*^+^(A and B). DIC images (C,F,I and C,F) illustrates external normal morphology in wild type and more pronounced tumorous phenotypes in homozygous *l(3)tb* larval brain and wing imaginal discs. Quantitative analysis showed increase in the number of mitotic positive cells in homozygous mutant larval brain lobes and wing imaginal discs as compared to wild type and difference was highly significant (IV). The images are projections of optical sections taken by confocal microscope, Scale bar 100μm (I, II) and 50 μm (III), Staining was done in triplicates with 10 brains and 15 wing imaginal discs in each group. Significant difference is represented as *** *P*≤0.005 using one-way ANOVA.

### The mutation *l(3)tb* alters expression of DE-cadherin in brain and wing imaginal discs and affects neuronal number and fasciculation

During the onset and progression of tumorigenesis, the mutation was observed to impart numerous perturbations in the developmental expression of essential molecules globally. DE-cadherin, a protein expressed in the adherens junctions besides being instrumental in neural development, was found to show altered expression in the homozygous mutant brain and wing imaginal discs in the late larval stages (Day 10) (**Figure 4D**). The wild type brain (Day 5; 115h ALH) however, showed strong expression in the neuropile and in the Outer and Inner Proliferating centres (OPC and IPC, respectively) of the brain hemispheres (**Figure 4A**). The central brain in *Drosophila* harbors the Mushroom Body (MB) and other neurons, which express the cell adhesion molecule Fasciclin II, which also labels the pioneering axonal tracts or fascicles in the neuropile and axonal projections in the ventral ganglion (**Figure 4B**). Mature mutant larvae showed disrupted arrangement of neurons of the MB and those of the central brain (**Figure 4E**). Expression analysis of the pan-neuronal marker, Elav showed a progressive decrease with increase in larval age and simultaneous maturation of the tumor (**Figure 6A, D and G**), implying progressive loss of neurons during the progression of tumorigenesis. Discs-large (Dlg), a septate junction marker, which is strongly expressed in the central brain as well as in the neurons and their projections emanating from the ventral nerve cord (VNC) (**Figure 6B**), showed enhanced expression in mature mutant larval brain (Day 10) as compared to early third instar larval brain (both wild type or mutant) but its pattern in the central brain and neuropile was lost (**Figure 6E and H**). The staining pattern is in agreement with results obtained from immunoblotting experiments (**Figure 6V**). Parallel to the disruption of DE-cadherin in the wing disc (**Figure 5E and I**), a progressive disruption of Armadillo, the *Drosophila* homologue of β-catenin and is associated with the E-cadherin junctions, was observed in the late/mature mutant larvae (**Figure 5F and J**). Closer analysis revealed that the tumorous wing disc showed enlarged cell sizes (**Figure 5Q**) as compared to the wild type (**Figure 5M**) or early stage tumor tissue and that these two proteins are co-expressed in the wing pouch (**Figure 5P**) but following the onset of tumorigenesis, the loss of armadillo occurs prior to that of DE-cadherin (**Figure 5T**).

**FIGURE 4.**
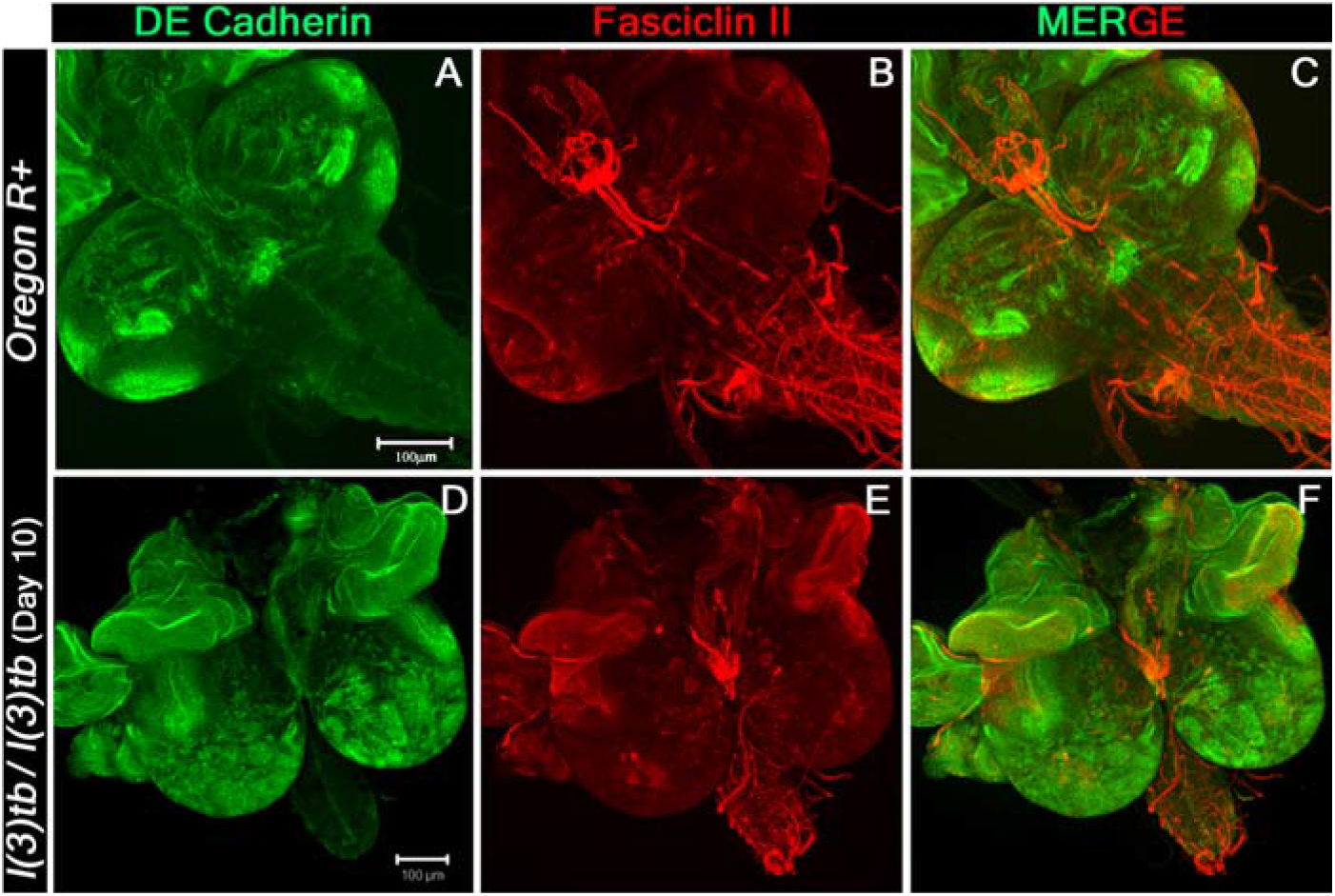
Confocal photomicrograph of 3^rd^ instar larval brain showing the distribution pattern of DE-cadherin (Green) and Fasciclin II (red). Strong expression of DE-Cad throughout the brain lobes seen in homozygous *l(3)tb* (D) whereas wild type shows specific regions (A). Fas II expression is also altered in homozygous *l(3)tb* (E) as compared to wild type (B). Scale bar 100 μm.

**FIGURE 5.**
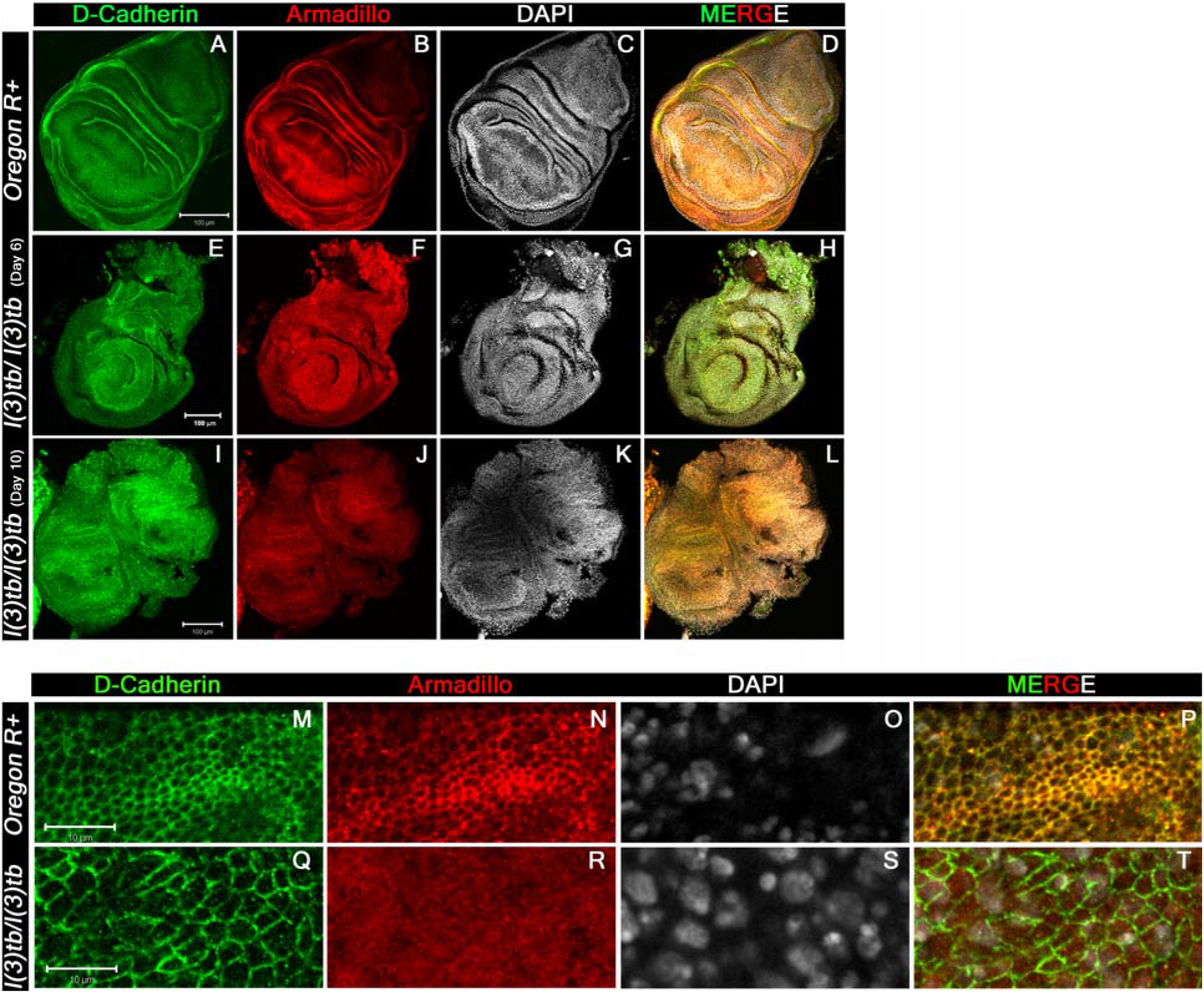
Confocal images of 3^rd^ instar larval wing imaginal discs immunolabeled to visualize the altered distribution pattern of cadherin-catenin complex proteins. Tumor caused in the homozygous *l(3)tb* mutant completely altered the distribution pattern of both, trans-membranous protein DE-cadherin (A, E, I, M, Q) and Armadillo (β-Catenin, B, F, J, N, R) adheren junctional proteins. Alteration of both proteins is more pronounced in the wing imaginal discs from mutant larva during extended larval life (I, J) than in the early wing imaginal disc (E, F) as compared to distinct pattern of DE-cadherin (A) and Armadillo (B) in the wild type wing imaginal discs. Armadillo is a binding partner of trans-membranous protein DE-cadherin having roles in cell adhesion and regulate tissue organization and morphogenesis. Merged images also substantiate the altered distribution of both junctional proteins in the homozygous mutant (H, L) as compared to the wild type (D) where co-localization is indicated by yellow pattern. Higher magnification of wing imaginal disc (pouch region) demonstrate altered distribution pattern of DE-cadherin (Q) and Armadillo (R) in homozygous *l(3)tb* mutant as compared to wild type (N, R). Increase in cell size seen in homozygous *l(3)tb* mutant (Q) as compared to wild type (M). Complete loss of Arm staining observed in homozygous *l(3)tb)(R)* whereas normal pattern seen in wild type wing disc (N). Chromatin size also altered in homozygous *l(3)tb* (S) as compared to wild type (O). Wild type shows clear co-localization of D-Cad and Arm (P), while there is complete loss of co-localization in homozygous *l(3)tb* wing imaginal discs (T). Scale bar represents 100 μm (A to L) and 10 μm (M to T).

### Eye-antennal discs and leg imaginal discs also show morphological and developmental anomalies in *l(3)tb* homozygous individuals

Global analysis of morphological aberrations in the mutant homozygotes showed that besides the tumorous brain and wing imaginal discs, eye-antennal discs and leg imaginal discs were also overgrown with a transparent appearance. Expression of Elav and Dlg in the eye-antennal discs revealed similarities to the developmental perturbations observed in the wing discs and brain. In the early third instar mutant larvae (Day 6), all photoreceptor cells showed expression of Elav, similar to the wild type tissue (**Figure 6J and N**). However, during advanced stages of larval tumorigenesis (Day 10), it dwindled eventually (**Figure 6R**). The Elav expressing cells which are posterior to the morphogenetic furrow co-express Dlg and demonstrate the typical ommatidial arrangement. In the mature mutant larvae however, the eye discs demonstrate significant deviations from the normal regular arrangement of ommatidia.

**FIGURE 6.**
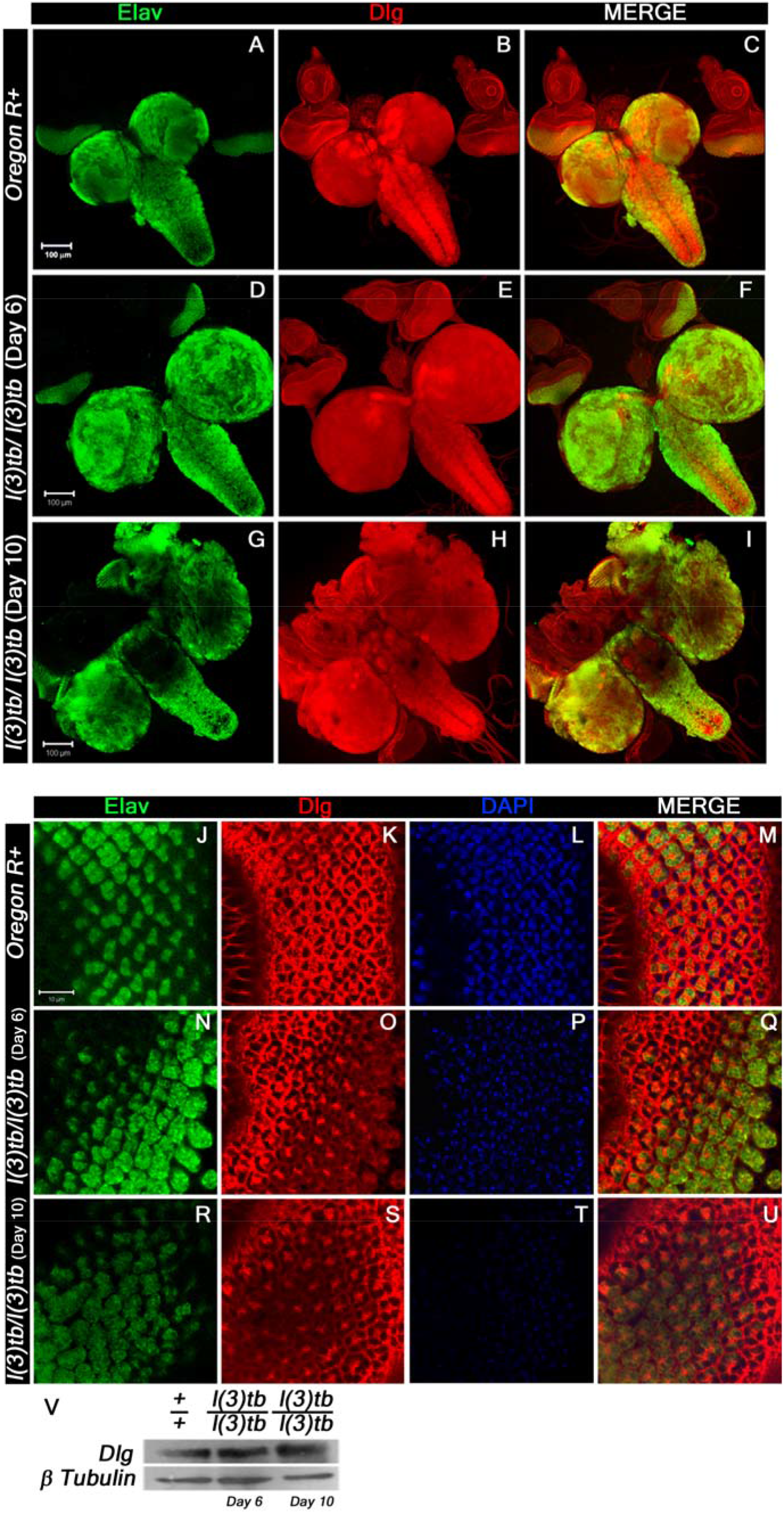
Confocal photomicrograph show loss of mature neurons and increase in junctional protein, Dlg, in delayed (Day 10) homozygous *l(3)tb*. 3^rd^ instar larval brain shows intense staining of Elav (green) in day 6 (D) of homozygous mutant later on show loss of staining in enlarged brain of day 10 (G), while the wild type brain (A) showed normal pattern of Elav staining. Dlg stained the ventral nerve chord and central brain in optic lobes of wild type (B), which is similar in day 6 of homozygous mutant brain (E) but in delayed larval brain, day 10, the pattern was altered (H). Scale shown is 100μm. Neuronal tissue from eye imaginal discs also display loss of neurons seen through Elav staining in day 10 (R) as compared to day 6 (N) in homozygous *l(3)tb* mutant as well as to wild type (J). Pattern of junctional protein, Dlg, in eye imaginal discs is also altered in day 10 (S) as compared to day 6 (O) and wild type (K). Counter stain with DAPI shows very weak intensity in day 10 (T) reflecting disintegrating chromatin as compared to day 6 (P) and wild type (L). Scale bar represents 10μm. Western blot for comparison of Dlg protein in wild-type (+/+), day 6 and day 10 *l(3)tb* mutant larval brain showed increased Dlg protein in homozygous mutant larval brain (V). β-tubulin has been used as an internal control.

The leg imaginal discs, which reside in close proximity to the brain and wing imaginal discs also show enlargement in size which increases with advancement and retention of larval stage. They show gradual disruption of normal expression of DE-cadherin and Armadillo, alike tumorous wing discs (see above), implying the mutation and subsequent tumor to affect developmental homoeostasis in adjacent tissues as well.

### Analysis of Meiotic Recombination and complementation mapping identify *l(3)tb* to be allelic to *DCP2*

The mutation *l(3)tb* was maintained with *TM6B* balancer, which established its localization on the third chromosome. Analysis of meiotic recombination frequencies of an unmapped mutation with known markers is a classical technique that has been routinely employed to identify its cytogenetic position. *Drosophila* has the advantage of having classical markers with well documented visible phenotypes for each chromosome, which aid in such mapping endeavours. In order to bring *l(3)tb* in a chromosome with such markers (*rucuca*), we allowed meiotic recombination to occur between *l(3)tb* and the 8 recessive markers present on the *rucuca* chromosome (**Table 2**). 113 recombinant males were observed and recombination frequencies were calculated in centiMorgan (cM). **Table 3** shows the recombination frequencies of each marker (locus) with the mutation *l(3)tb*. Preliminary analysis suggested that *l(3)tb* was close to *thread (th)* with minimum recombination events between the two loci (2.65%). Further analysis of recombination events between *h-l(3)tb* [17.78%], *st-l(3)tb*[1.23%] and *cu-l(3)tb*[8.29%] (**Table 4**) and comparing with the positions of each of the markers, the mutation was estimated to be located left of *thread* (43.2 cM; band 72D1) between 41.71 cM–42.77 cM, *i.e*., in the cytological position 71F4-F5.

**TABLE 2.**
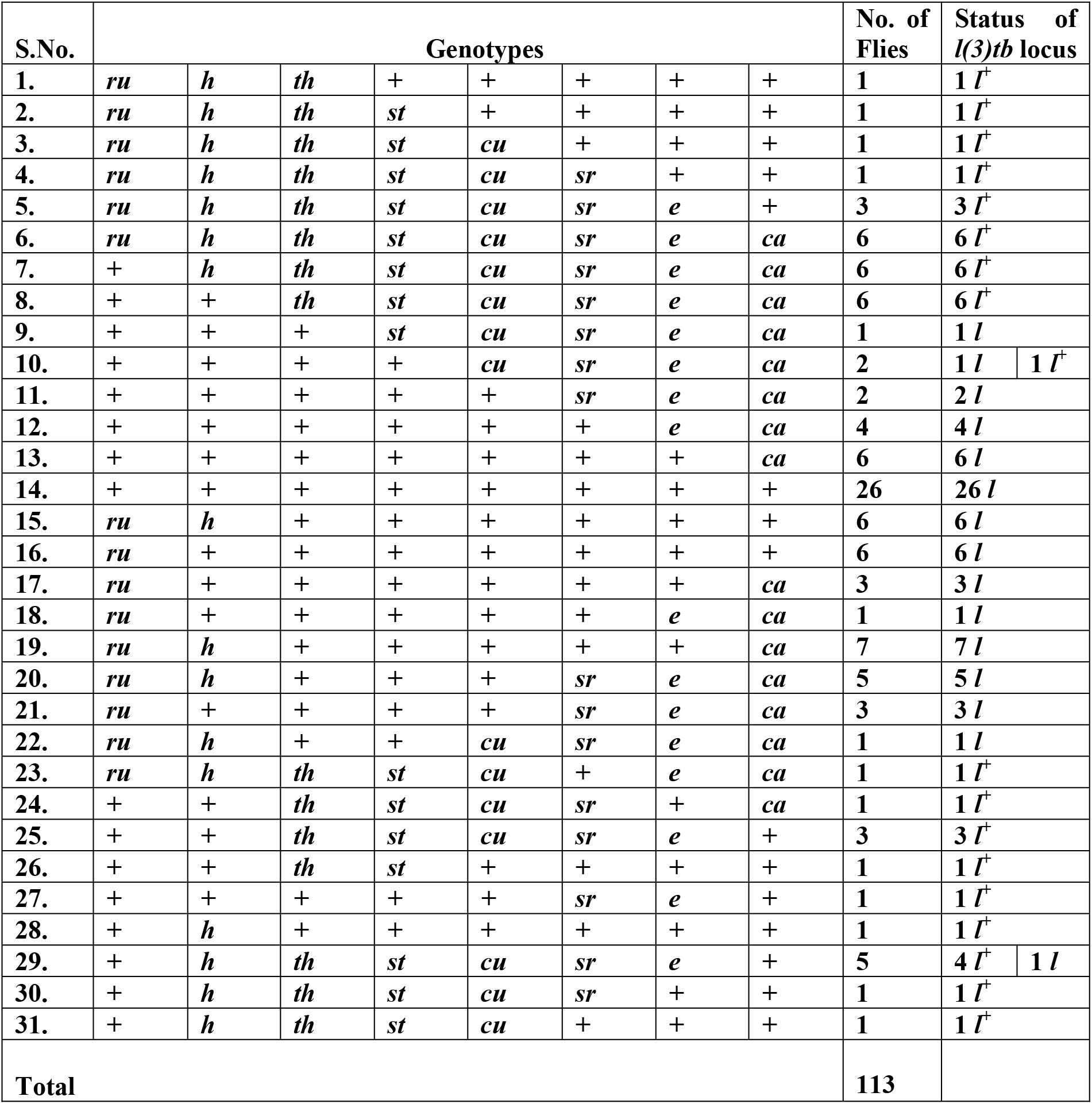
Rearranged genotypes of 113 males after various recombination events between all the eight visible markers of *rucuca* chromosome

**TABLE 3.** Recombination frequencies (RF) between various recessive markers on *rucuca* chromosomes (*roughoid, hairy, thread, scarlet, curled, stripe, ebony*, and *claret*) and *l(3)tb*

| Sl. No. | Association of marker with *l(3)tb* | Flies |  |  |  | Recombination frequency (RF) |
| --- | --- | --- | --- | --- | --- | --- |
|  |  | Parental (P) |  | Recombinant (R) |  |  |
| | | Genotype | No. of flies | Genotype | No. of flies | $\frac{R}{P+R} \times 100$ |
| 1. | *ru - l* | *ru*<sup>+</sup> *l*<br>*ru* *l*<sup>+</sup> | 18<br>} 62<br>44 | *ru*<sup>+</sup> *l*<sup>+</sup><br>*ru* *l* | 18<br>} 51<br>33 | 45.13 |
| 2. | *h - l* | *h*<sup>+</sup> *l*<br>*h* *l*<sup>+</sup> | 54<br>} 79<br>25 | *h*<sup>+</sup> *l*<sup>+</sup><br>*h* *l* | 12<br>} 34<br>22 | 30.08 |
| 3. | *th - l* | *th*<sup>+</sup> *l*<br>*th* *l*<sup>+</sup> | 74<br>} 110<br>36 | *th*<sup>+</sup> *l*<sup>+</sup><br>*th* *l* | 1<br>} 3<br>2 | 2.65 |
| 4. | *st - l* | *st*<sup>+</sup> *l*<br>*st* *l*<sup>+</sup> | 73<br>} 108<br>35 | *st*<sup>+</sup> *l*<sup>+</sup><br>*st* *l* | 2<br>} 5<br>3 | 4.42 |
| 5. | *cu - l* | *cu*<sup>+</sup> *l*<br>*cu* *l*<sup>+</sup> | 71<br>} 105<br>34 | *cu*<sup>+</sup> *l*<sup>+</sup><br>*cu* *l* | 3<br>} 8<br>5 | 7.07 |
| 6. | *sr - l* | *sr*<sup>+</sup> *l*<br>*sr* *l*<sup>+</sup> | 62<br>} 94<br>32 | *sr*<sup>+</sup> *l*<sup>+</sup><br>*sr* *l* | 4<br>} 19<br>15 | 16.8 |
| 7. | *e - l* | *e*<sup>+</sup> *l*<br>*e* *l*<sup>+</sup> | 56<br>} 86<br>30 | *e*<sup>+</sup> *l*<sup>+</sup><br>*e* *l* | 7<br>} 27<br>20 | 23.89 |
| 8. | *ca - l* | *ca*<sup>+</sup> *l*<br>*ca* *l*<sup>+</sup> | 42<br>} 74<br>32 | *ca*<sup>+</sup> *l*<sup>+</sup><br>*ca* *l* | 16<br>} 49<br>33 | 43.3 |

**TABLE 4.** Recombination events between *h-l, st-l* and *cu-l*

| S.No. | Association<br>of marker<br>with *l(3)tb* | Flies | | | | Recombination<br>frequency<br>$\frac{R}{P+R} \times 100$ |
| --- | --- | --- | --- | --- | --- | --- |
|  |  | Parentals (P) |  | Recombinants (R) |  |  |
|  |  | Genotype | No. of flies | Genotype | No. of flies |  |
| 1. | *h - l* | *h<sup>+</sup> l* | 89 | *h<sup>+</sup> l<sup>+</sup>* | 18 | 17.78 |
|  |  | *h l<sup>+</sup>* | 96 | *h l* | 22 |  |
|  |  |  | 185 |  | 40 |  |
| 2. | *st - l* | *st<sup>+</sup> l* | 252 | *th<sup>+</sup> l<sup>+</sup>* | 2 | 1.23 |
|  |  | *st l<sup>+</sup>* | 238 | *th l* | 4 |  |
|  |  |  | 480 |  | 6 |  |
| 3. | *cu - l* | *cu<sup>+</sup> l* | 269 | *ru<sup>+</sup> l<sup>+</sup>* | 22 | 8.29 |
|  |  | *cu l<sup>+</sup>* | 283 | *ru l* | 28 |  |
|  |  |  | 552 |  | 50 |  |

Complementation analysis with molecularly defined Drosdel and Exelixis deficiency lines (N=85), spanning the entire chromosome 3, identified four lines which failed to complement the mutation, *viz., Df(3L)BSC774, Df(3L)BSC575, Df(3L)BSC845* and *Df(3L)RM95*, which was generated in the lab using progenitor RS stocks. Trans-heterozygotes *l(3)tb/Df(3L)BSC575* were pupal lethal and the dying non-tubby larvae showed phenotypes similar to *l(3)tb* homozygotes, suggesting the mutation to reside between 71F1 and 72A1 on the left arm of chromosome 3. Further analysis using six deletion lines belonging to the above region (71F1–72A2) identified the mutation to reside between 71F4 to 71F5, which strangely is a gene desert region. Complementation analyses performed with lethal insertion alleles (N=26) of genes residing proximal or distal to 71F4-F5 identified two lethal P-element insertion alleles of *DCP2* (mRNA decapping protein 2; CG6169), *viz., P{GT1}Dcp2^BG01766^* and *PBac{RB}Dcp2^e00034^*, which failed to complement the mutation *l(3)tb* (**Figure 7A and C**) as well as those deletions which had failed to complement *l(3)tb*, implying the mutation to be allelic to *DCP2* (72A1).

**FIGURE 7.**
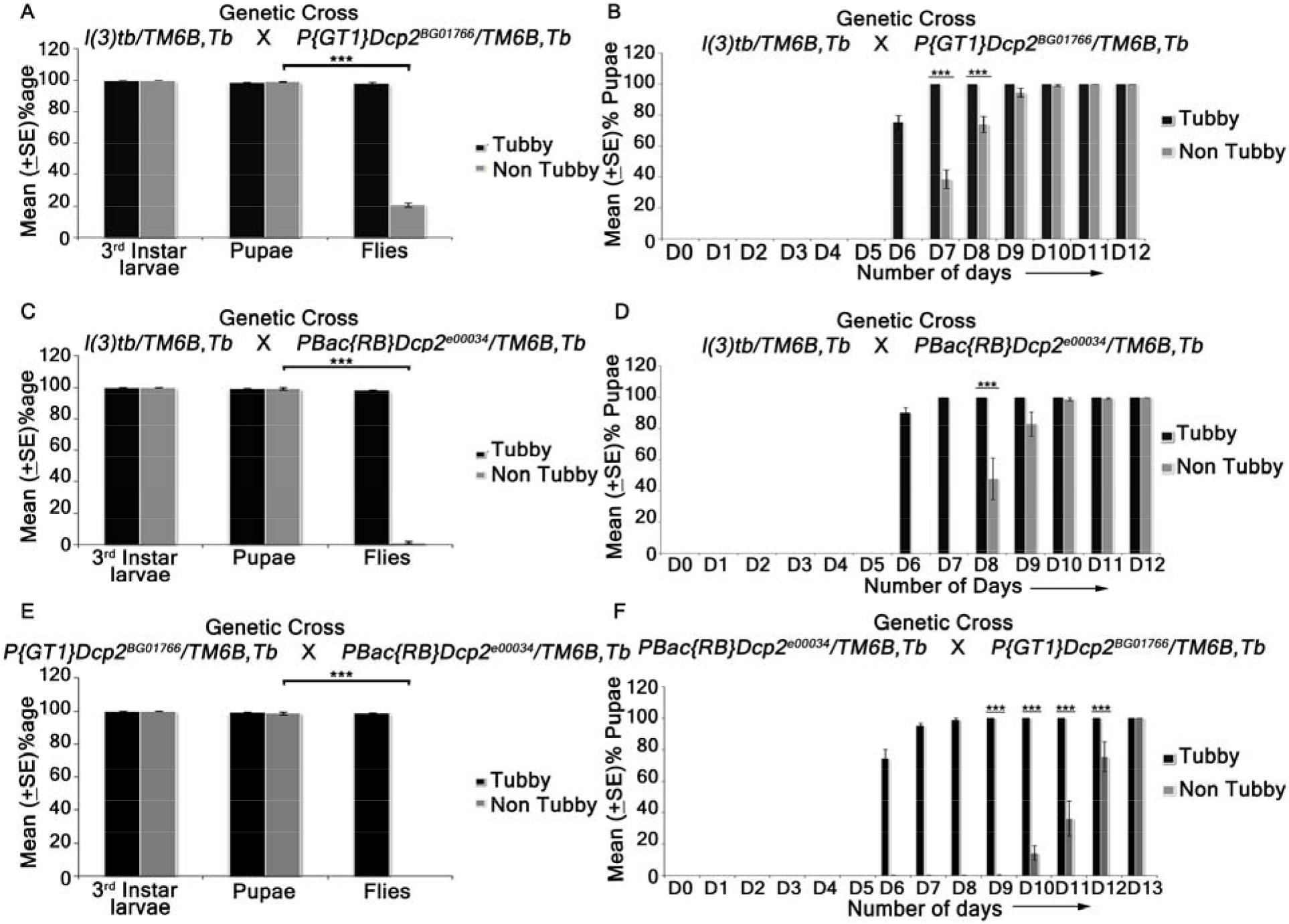
Viability assay performed on various hetero-allelic combinations between alleles of gene *DCP2* and the mutation in *l(3)tb*. Homozygous *l(3)tb* exhibited larval as well as pupal lethality. 69% of homozygous larvae pupated whereas no fly eclosed from the pupae. *l(3)tb* trans-heterozygous with *P{GT1}DCP2^BG01766^* showed only 18.4% fly eclosed (A). *l(3)tb/PBac{RB}DCP2^e00034^* trans-heterozygote (C) causes 100% lethality at pupal stage. Trans-allelic combination *P{GT1}DCP2^BG01766^//PBac{RB}DCP2^e00034^* (E) also exhibited 100% pupal lethality. Developmental delay seen in trans-heterozygotes *l(3)tb /P{GT1}DCP2^BG01766^* (B) and *l(3)tb/PBac{RB}DCP2^e00034^* (D) as in homozygous *l(3)tb*. Progeny from heterozygous for both the alleles of *DCP2* gene, *PBac{RB}DCP2^e00034^ /P{GT1}DCP2^BG01766^* (F) also exhibited developmental delay. *** indicates p<0.005.

### Trans-heterozygotes of *DCP2* mutants and *l(3)tb* show developmental delay, tumorous larval brain with elevated neuroblast numbers, larval/pupal lethality and developmental defects in escapee flies

Trans-heterozygotes of *l(3)tb* with either allele of *DCP2, viz., P{GT1}Dcp2^BG01766^* and *PBac{RB}Dcp2^e00034^*, showed developmental delay. In either case, trans-heterozygous third instar larvae showed persistence of larval stage till Day 10 ALH (**Figure 7B and D**), and show tumorous phenotypes of brain and wing imaginal discs (data not shown), similar to the *l(3)tb* homozygotes. Expression pattern of Deadpan (Dpn), a marker for neuroblasts show increased number of neuroblasts in the larval brain of the trans-heterozygotes as well as *l(3)tb* homozygotes (**Figure 8F, K and P**). Also, the trans-heterozygous progeny showed a higher mitotic index as compared to the wild type progeny, similar to the *l(3)tb* homozygotes (**Figure 8G, L and P**). While *PBac{RB}Dcp2^e00034^/l(3)tb* was found to be 100% pupal lethal, *P{GT1}Dcp2^BG01766^/l(3)tb* was only 81.6% lethal (**Figure 7A and B**), with the rest 18.4% pupae eclosing as flies. However, the escapee flies showed several developmental abnormalities, *viz*., defects in wing (9.5%), thorax closure (3.2%), loss of abdominal para-segments and abdominal bristles (3.2%), and presence of melanotic patches (22.2%), leg defects (41.3%) or eclosion defects (12.7%). Analysis of compound eyes in these escapees revealed complete loss of regular arrangement of ommatidia and ommatidial bristles. Abnormal external genitalia were also observed in the male escapees. Subsequent analysis of fertility showed that the trans-heterozygous escapee flies had compromised fertility with only 40% of the males and 21.7% of the females being fertile (**Table 5**).

**FIGURE 8.**
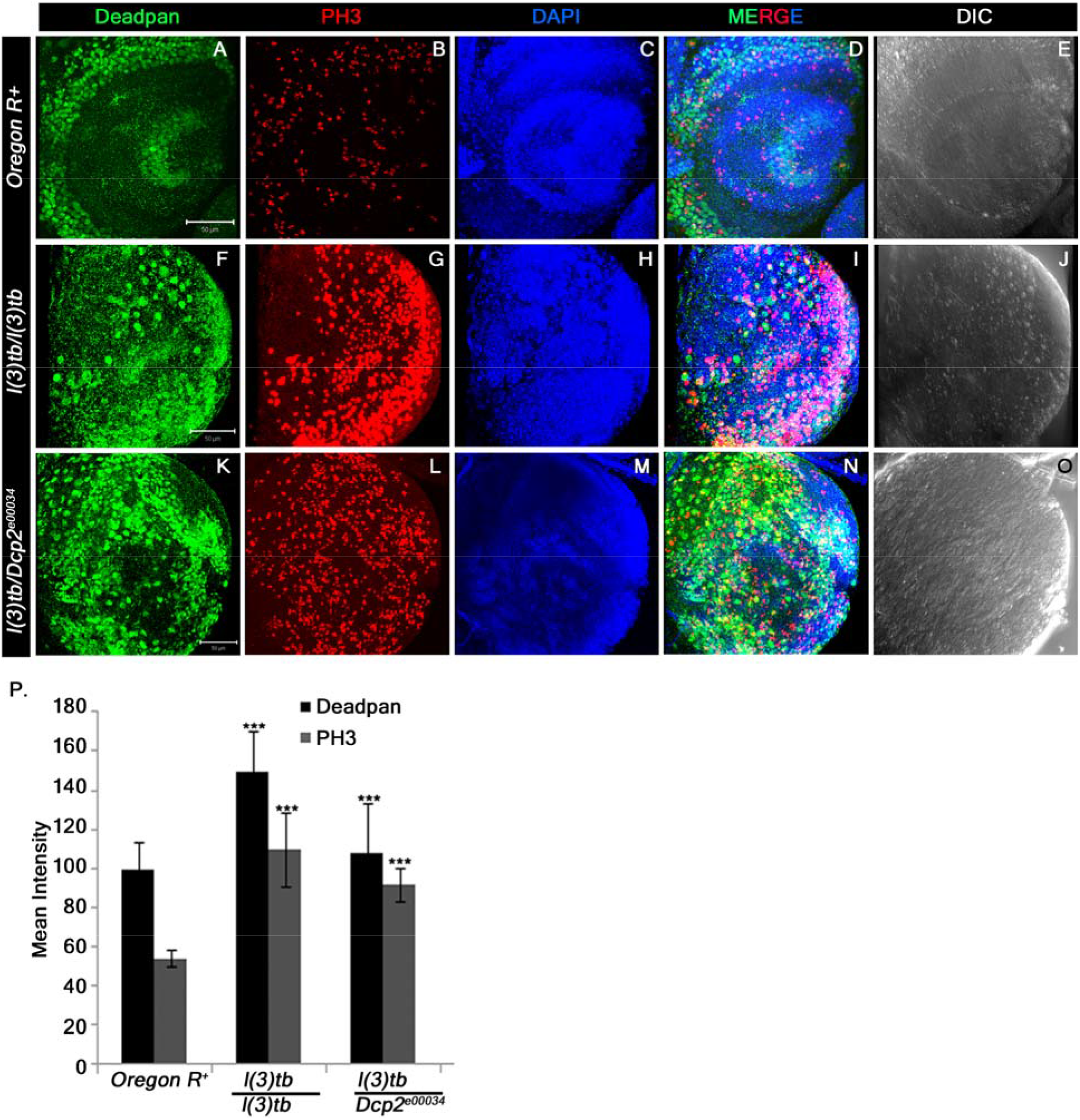
Heterozygous combination of *l(3)tb* with *DCP2^e00034^* allele resulted in to significant increase in the number of neuroblasts and mitotically active cells. Confocal projection sections showing immunolocalisation of Deadpan, a neuroblast marker (Green, A, F, K) for picking neuroblasts and phosphohistone 3 (PH3, red, B, G, L) marking the mitotic cells are shown. Enhanced neuroblast population in homozygous mutant (F) and in heterozygous *l(3)tb* with *DCP2^e00034^* allele (K) Similarly, increased number of mitotic cells (PH3 positive) also occurred in heterozygous *l(3)tb* with *DCP2^e00034^* allele (L), similar to homozygous *l(3)tb* mutant (G). NBs and mitotic positive cells are quantified (P) and the differences are statistically significant when compared with wild type. *** *P* >0.005. Scale bar indicates 50μm.

**TABLE 5.**
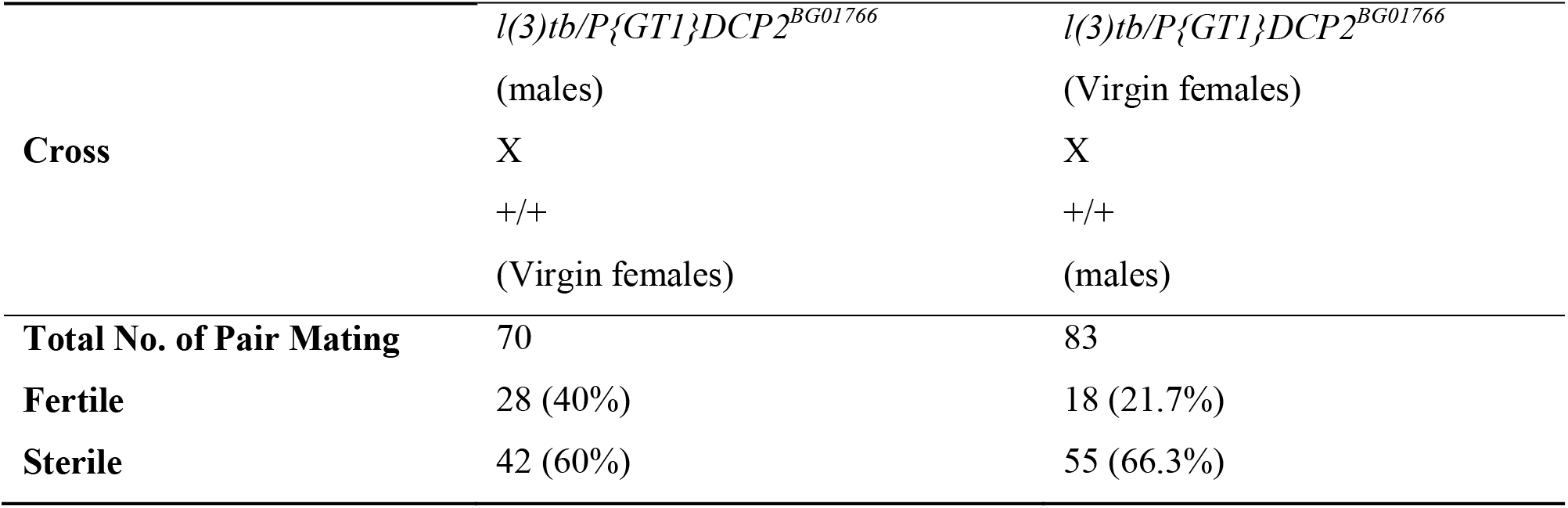
Fertility assay of trans-heterozygotes *P{GT1}DCP2^BG01766^/l(3)tb* demonstrating male and female sterility

The similarity in the pattern of development and the defects associated with it between the *l(3)tb* trans-heterozygotes and homozygotes provide a strong genetic proof of allelism between *l(3)tb* and *DCP2*.

### A single nucleotide mutation in the promoter region affects transcription of *DCP2* in *l(3)tb* mutants

To identify the sequence alterations in *DCP2* in the *l(3)tb* homozygotes, 28 pairs of overlapping primers were designed, spanning the entire gene (7.689 kb). **Figure 9** shows the four mutations identified, *viz*.,**G**(3L:15819202)**A**, G(3L: 15819384)**A**, C(3L:15819446)**T** and **C**(3L:15819691)**A**, out of which **C**(3L:15819691)**A** resides in the promoter DCP2_1 (Eukaryotic Promoter Database; EPD, SIB) (**Figure 9E**). *DCP2* codes for four transcripts, *viz*., DCP2-RA, RB, RD and RE. All transcripts differ in their 5’ and 3’ UTRs but have conserved exon sequences. Analysis of gene expression using primers designed to amplify the exon common to all the transcripts showed absence of amplification (**Figure 9F**), implying the mutations identified above to affect gene transcription.

**FIGURE 9.**
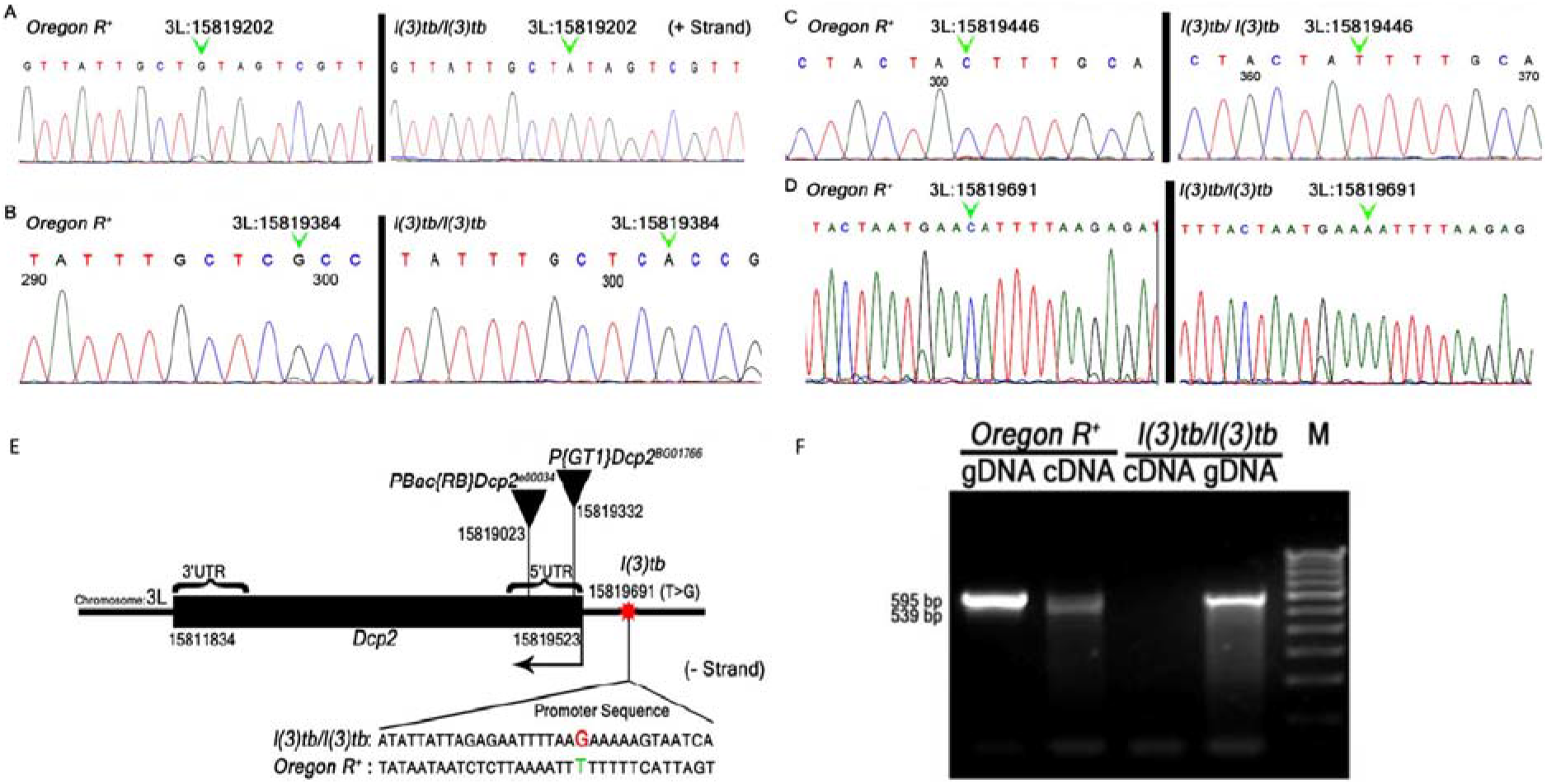
Mutations found in the 5’ UTR and promoter of *DCP2* in genomic DNA of *l(3)tb* homozygotes. Chromatograms show single nucleotide changes at positions **3L:15819202(A>G); 3L:15819384(A>G): 3L:15819446(A>C)** on plus strand (A-D). Three mutations lie in the 5’UTR of *DCP2* and one transversion **3L:15819691(T>G)** lie in the promoter of *DCP2*. All nucleotides are numbered according to FB2013_05 *Dmel* Release 5.53 (FlyBase). A schematic representation of *DCP2* depicting the insertion of the P-elements in the two insertion alleles, *viz., DCP2^e00034^* and *DCP2^BG01766^* is shown in E. Note the nucleotide change in the promoter in the *l(3)tb* mutant. Analysis of *DCP2* expression shows absence of DCP2 transcripts in homozygous *l(3)tb* mutant (F). Note that the primer pair generates different sized amplicons for genomic and cDNA. Although the genomic region coding for the transcript is present in the mutant genome, it fails to code for the same.

### Global overexpression of *DCP2* rescues mutant phenotypes associated with *l(3)tb*

Global over-expression of *DCP2* using ubiquitous GAL4 drivers (*Act5C-GAL4* or *Tub-GAL4*) in the mutant homozygous *l(3)tb* individuals rescued the larval and pupal lethality. **Table 6** shows the genotype and fate of the progeny as scored from the rescue experiment. As can be seen, for over-expression of *DCP2* using *Act5C-GAL4*, out of 35.1% (N=155) non-tubby progeny *(l(3)tb* homozygous background), *i.e., Act5C-GAL4/CyO* or *Sp; l(3)tb:UAS-DCP2/l(3)tb*, 21.3% (N=94) and 13.8% (N=61) segregated as curly (*Act5C-GAL4/CyO; l(3)tb:UAS-DCP2/l(3)tb*) and non-curly (or with sternopleural bristles: *Act5C-GAL4/Sp; l(3)tb:UAS-DCP2/l(3)tb*), respectively. Similarly, while over-expressing using *Tub-GAL4*, we obtained 37% (N=166) non-tubby progeny, *i.e., UAS-DCP2/CyO* or *Sp; l(3)tb:Tub-GAL4/l(3)tb*, out of which, 17.2% (N=77) were curly (*UAS-DCP2/CyO; l(3)tb:Tub-GAL4/l(3)tb*) while 19.8% (N=89) were non-curly (*UAS-DCP2/Sp; l(3)tb:Tub-GAL4/l(3)tb*).

**Table 6.** Global overexpression of *DCP2* rescues the mutant phenotypes exhibited by *l(3)tb*

| Genetic Crosses |  | *Act5C-GAL4/CyO*;<br>+/+ |  | +/+; *Tub-GAL4/TM6B* |  | *Act5C-GAL4/CyO*;<br>*l(3)tb/TM6B* |  | *UAS-DCP2/CyO*;<br>*l(3)tb/TM6B* |  |
| --- | --- | --- | --- | --- | --- | --- | --- | --- | --- |
|  |  | X |  | X |  | X |  | X |  |
|  |  | *Act5C GAL4/CyO*;<br>+/+ |  | +/+; *Tub GAL4/TM6B* |  | *Sp/CyO*; *l(3)tb: UAS-DCP2/TM6B* |  | *Sp/CyO*; *l(3)tb: Tub GAL4/TM6B* |  |
|  |  | Homozygotes die as embryos or early larvae |  | Homozygotes die as embryos or early larvae |  | *CyO* and *TM6B* homozygotes die as embryos or early larvae |  | *CyO* and *TM6B* homozygotes die as embryos or early larvae |  |
| 01. | Eggs | **750** |  | **790** |  | **1050** |  | **1245** |  |
| 02. | Unfertilised Eggs | 39 | 5.2% | 37 | 4.7% | 68 | 6.5% | 86 | 6.9% |
| 03. | Fertilised Eggs | 711 | 94.8% | 753 | 95.3% | 982 | 93.5% | 1159 | 93.1% |
| 04. | Dead Embryos | 304 | 42.8% | 357 | 47.4% | 434 | 44.2% | 525 | 45.3% |
| 05. | Dead 1<sup>st</sup> and 2<sup>nd</sup> instar Larvae | 8 | 1.12% | 19 | 2.52% | 34 | 3.46% | 57 | 4.92% |
| 06. | Dead 3<sup>rd</sup> instar larvae | 2 | 0.28% | 5 | 0.66% | 72 | 7.33% | 128 | 11.04 % |
| 07. | Pupae | **397** | 55.8% | **372** | 49.4% | **442** | 45.0% | **449** | 38.7% |
| 08. | Dead Pupae | 17 | 4.3% | 11 | 2.9% | 21 | 4.8% | 23 | 5.1% |
| 09. | Eclosion following over-expression of *DCP2* in homozygous *l(3)tb* background | - | - | - | - | *Act5C-GAL4/CyO*;<br>*l(3)tb: UAS-DCP2/l(3)tb* |  | *UAS-DCP2/CyO*;<br>*l(3)tb: Tub-GAL4/l(3)tb* |  |
|  |  | - | - | - | - | 94 | **21.3%** | 77 | **17.2%** |
|  |  | - | - | - | - | *Act5C-GAL4/Sp*;<br>*l(3)tb: UAS-DCP2/l(3)tb* |  | *UAS-DCP2/Sp*;<br>*l(3)tb: Tub-GAL4/l(3)tb* |  |
|  |  | - | - | - | - | 61 | **13.8%** | 89 | **19.8%** |

In both the cases of overexpression, all non-tubby progeny pupated, devoid of any developmental anomalies reminiscent of *l(3)tb* mutation and emerged as flies. Thus, the rescue of the mutant phenotypes observed in *l(3)tb* homozygotes by global overexpression of *DCP2* iteratively substantiates the fact the *l(3)tb* is an allele of *DCP2* and that the tumor is caused solely owing to the loss of expression of *DCP2*.

## Discussion

Mutants provide an excellent platform for exploration of gene function. In the present communication, we have mapped and characterized the phenotype of a novel mutation, *lethal(3)tumorous brain* [*l(3)tb*], which was found to allelic to *DCP2*, the mRNA decapping protein 2 in *Drosophila melanogaster*. Like other well established tumor suppressor mutants in *Drosophila, viz., lethal(2) giant larvae* [*l(2)gl*], discs-large [Dlg] and *Scribble* [Scrib], *l(3)tb* homozygous embryos progress into and through the larval stages, grow normally until the third instar larval stage, undergo an extended larval life unlike normal individuals, pupate and then die. During the extended larval period, the larvae become bloated, transparent and the proliferating imaginal disc epithelia and nervous system appear dramatically aberrant. All these phenotypes are shared by most of the tumor suppressor mutants in *Drosophila*. The mitotic chromosomes in *l(3)tb* mutant show extended chromosomes albeit without any numerical anomaly. These secondary chromosomal changes also occur during the course of mammalian tumor progression and have some role in conferring metastatic potential to the tumor tissue (Yunis, 1983). In many tumors, cell cycle is not dramatically altered but cells fail to respond to arrest cues and cause over-proliferation. In *Drosophila*, the overall size of the tissue rather than number of cells *per se* is regulated precisely (Johnston and Gallant, 2002). Woods and Bryant reported (Woods and Bryant, 1989) that in neoplastic tumors, cell sizes are smaller initially but maintain their proliferative state during the prolonged larval phase which grossly alters the morphology of the imaginal discs. The *l(3)tb* mutants, both homozygotes and heterozygotes with *DCP2* mutants showed a similar behavior along with a high mitotic index as evidenced by PH3 staining, and thus the overgrowth in the discs can be ascribed to failure of the cells to exit the precisely coordinated developmental cell cycles. Although the number of neuroblasts in these mutants was pronounced, as revealed by the expression of Deadpan and PH3, the expression of Elav showed that the number of neurons was reduced, which may be attributed to the arrest of terminal differentiation from neuroblasts to neurons and/or glia, similar to that observed in *brat* mutants (Bello et al, 2006; Betschinger et al, 2006). Elav is a transcription factor and regulates the expression of certain neuronal genes, one of the target genes being *Armadillo*, the *Drosophila* homologue of β-catenin. Armadillo is expressed in the adherens junctions along with DE-cadherin, while Dlg is expressed at the septate junctions. The altered profile of Armadillo, DE-cadherin and Dlg and the simultaneous loss of Elav in the later larval stages in the *l(3)tb* mutants may be due to disassembly of the adherens junctions (Cox et al, 1996) thereby affecting cell-cell adhesion between glial and neuronal bodies in the brain and in the over-proliferative cells of the wing imaginal disc. There exists an inverse relation between E-cadherin function and tumor progression (Derksen et al, 2006) as E-cadherin plausibly regulates β-catenin signaling in the Wnt pathway with a potential to inhibit mitogenic signaling through growth factor receptors. This facet of *l(3)tb* tumors is similar to the simultaneous loss of E-cadherin and β-catenin observed in advanced stages in majority of mammalian tumors (Christofori and Semb, 1999; Weinberg and Hanahan, 2000).

Identification of cytogenetic location of mutations is best performed by calculation of recombination frequencies between the unknown mutation and the position of known loci (Sturtevant, 1913), and hence, recombination mapping, complementation analyses with regional deficiencies and duplications (Bridges, 1917 & 1919; Muller, 1935; Cook et al, 2012) was performed which revealed that the mutation resides close to the *thread* locus (43.2 cM, 72D1) between 71F4-F5, a gene desert area on the left arm of chromosome 3. Further complementation analysis with genes downstream to the region identified the failure of *DCP2* (CG6169) mutants, *viz., DCP2^BG01766^* and *DCP2^e00034^*, to complement the mutation as well as show phenotypes similar to *l(3)tb* homozygotes, thereby establishing the allelic relationship of *l(3)tb* to *DCP2. DCP2* is located at 72A1, which is adjacent to the region identified, *i.e*., 71F4-F5, and is thus in agreement with the results obtained from recombination and/or deficiency mapping.

The eukaryotic promoter database identifies 2 promoters for *DCP2*, DCP2_1 and DCP2_2. Sequencing analysis revealed a single transversion **C(15819691)A** in the promoter, DCP2_1, which is expected to affect its transcription and the subsequent expression of the gene, which is clearly observed in expression analysis of *DCP2* in *l(3)tb* homozygotes. The alleles themselves are embryonic lethal when homozygous, but trans-heterozygotes, *l(3)tb /DCP2^e00034^* and *l(3)tb/DCP2^BG01766^* show morphological, physical and physiological phenotypes similar to *l(3)tb* homozygotes. These trans-heterozygotes were tumorous with prolonged larval life, show increase in neuroblast population, elevated mitoses, and pupal lethality, all of which are exemplified by *l(3)tb* homozygotes. Parallely, all these phenotypes were rescued by global over-expression of *DCP2* in the *l(3)tb* homozygous mutant background, thereby validating the allelism between *DCP2* and the mutation *l(3)tb* and hence, we refer to the mutant line as an established *loss-of-function* allele of *DCP2* with defined genetic and molecular bases of allelism, *DCP2^l(3)tb^*. The cognate function of *DCP2* is removal of the 7-methylguanosine cap from the 5’ end of mRNAs, exposing them to the exonuclease, XRNI (Pacman) for degradation. In *Drosophila*, DCP2 is the only decapping enzyme present and thus is extremely important for a number of growth processes throughout development. In other organisms as well, it is extremely conserved and has fundamentally important roles in development (Xu et al, 2006; Ma et al, 2013), DNA replication (Mullen and Marzluff, 2008; Schmidt et al, 2011), stress response (Hilgers et al, 2006; Xu and Chua, 2012), synapse plasticity (Hillebrand et al, 2010), retrotransposition (Dutko et al, 2010) and viral replication (Hopkins et al, 2013). In *Arabidopsis, DCP2* loss-of-function alleles show accumulation of capped mRNA intermediates, lethality of seedlings and defects in post-embryonic development, with no leaves, stunted roots with swollen root hairs, chlorotic cotyledons and swollen hypocotyls (Goeres et al, 2007; Iwasaki et al, 2007; Xu et al, 2006). In humans as well, 5q21-22, the region harboring *DCP2* is frequently deleted in lung cancer (Hosoe et al, 1994; Mendes-da-Silva et al, 2000), colorectal cancer (Ashton-Richardt et al, 1989; Delattre et al, 1989) and oral squamous cell carcinoma (Mao et al, 1998). Cancer progression is associated with hyperactivated rRNA biogenesis (Chem et al 2011, Hein et al 2013) owing to the increased size and number of nucleoli (Pianese, 1896). Recently, Gaviraghi and coworkers (2018) have provided a plausible mechanism of regulation of aberrant rRNA biogenesis and/or maturation wherein the tumor suppressor PNRC1 recruits the decapping complex (DCP1/2) to the nucleolus and modulates the synthesis and maturation of U3 and U8 rRNAs. Hence, *DCP2* has an unexplored role in development and/or tumorigenesis throughout phyla, which needs to be investigated. Ren and coworkers reported that *DCP2* is expressed in the adult *Drosophila* brain (Ren et al, 2012). The pronounced defects observed in *l(3)tb* homozygotes also pertain to the brain and wing disc. Hence, the loss of *DCP2* in these tissues may affect the developmental homoeostasis existing in the gene expression network and may lead to tumorigenesis. We speculate that *DCP2* is potent enough to regulate the highly dynamic gene expression modules by virtue of its pioneering role in the mRNA decay pathways. Since the physiology of an organism is tightly regulated by the optimized titres of gene expression programs, a global loss of *DCP2* may lead to perturbed mRNA titres which in turn may alter the cellular response to such dismal conditions and eventually lead to drastic physiological disorders such as tumorigenesis. Although we are unsure of the exact mechanism(s) by which loss of *DCP2* leads to tumorigenesis, our findings in the allele, *DCP2*^l(3)tb^, propose an absolutely novel role of *DCP2* in tumorigenesis and identify *DCP2* as a candidate for future explorations of tumorigenesis.

---

**FIGURE S1.**
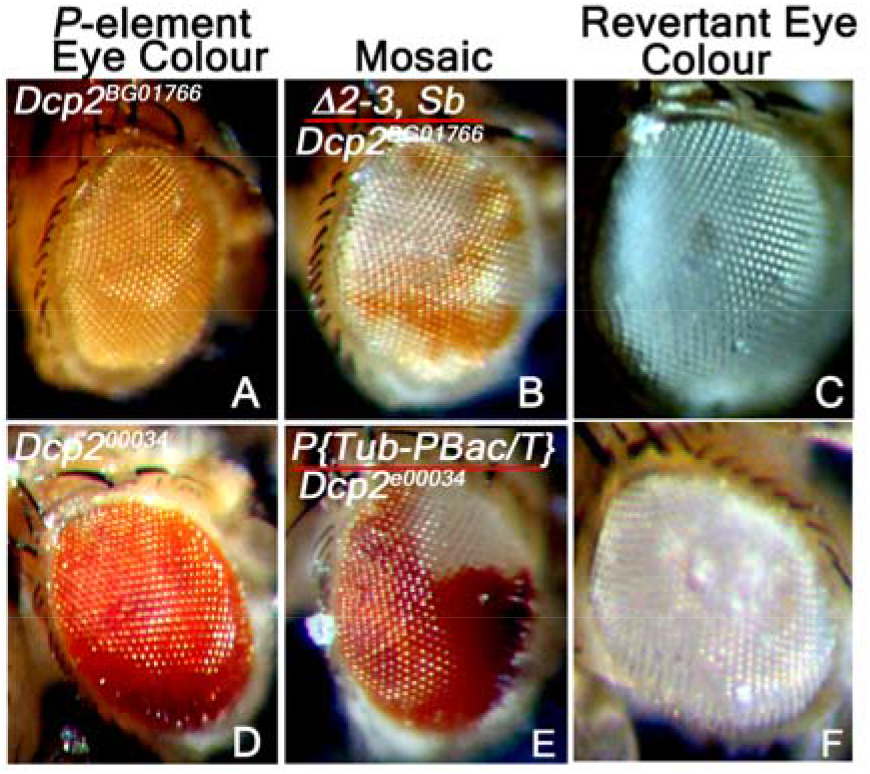
Reversion analysis by the excision of *piggyBac* transposon in *DCP2^e00034^* with the help of *piggyBac* specific transposase source, *CyO, P{Tub-Pbac}2/Wg^SP-1^* and similarly by the excision of *P*-element in *DCP2^BG01766^* strain using *Δ2-3,Sb/TM6B, Tb^1^, Hu, e^1^* transposase source as ‘jumpstarter stock’. DCP2 revertant white eyed F2 flies were crossed to *l(3)tb* and lethal progenies scored.

**FIGURE S2.**
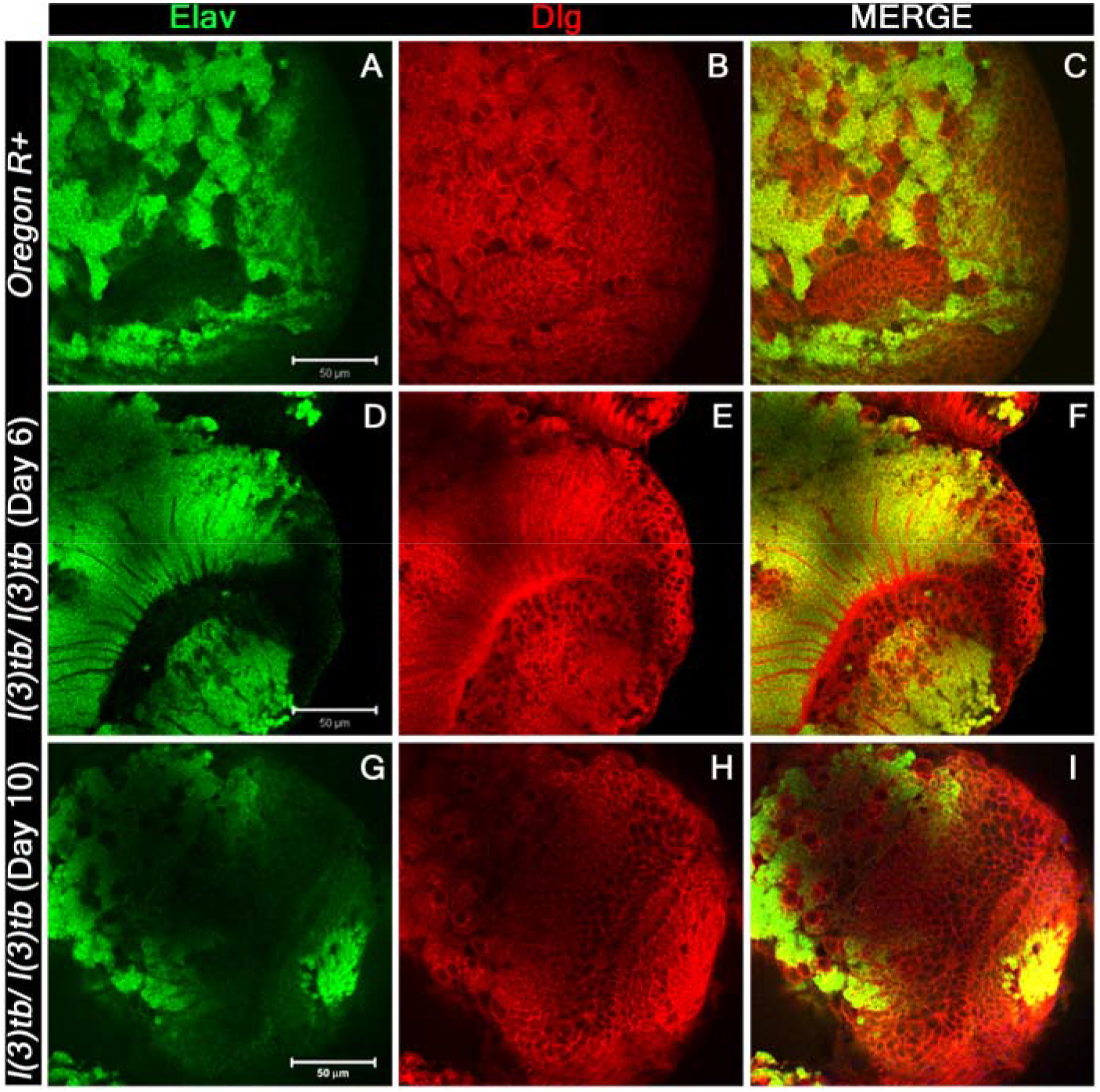
Confocal images of 3^rd^ instar larval brain lobe to visualize neurons through Elav (neuronal marker, green) co-labelled with septate junction marker Discs large, Dlg, (red). Confocal images of 3^rd^ instar larval brain to show neurons through Elav (green) and co-labelled with Dlg (red) at higher magnification. This also showed intense staining of Elav in day 6 (D) of homozygous mutant larval brain lobe which showed loss of staining in enlarged brain lobe of day 10 (G), while the wild type brain lobe (A) showed normal pattern of Elav staining. Dlg stained both lial and neuronal cells and showed distinct pattern in optic lobes of wild type larval brain (B), which is similar in day 6 of homozygous mutant brain (E) but in delayed enlarged larval brain lobe, day 10, the pattern was altered (H). Scale shown is 50μm.

**FIGURE S3.**
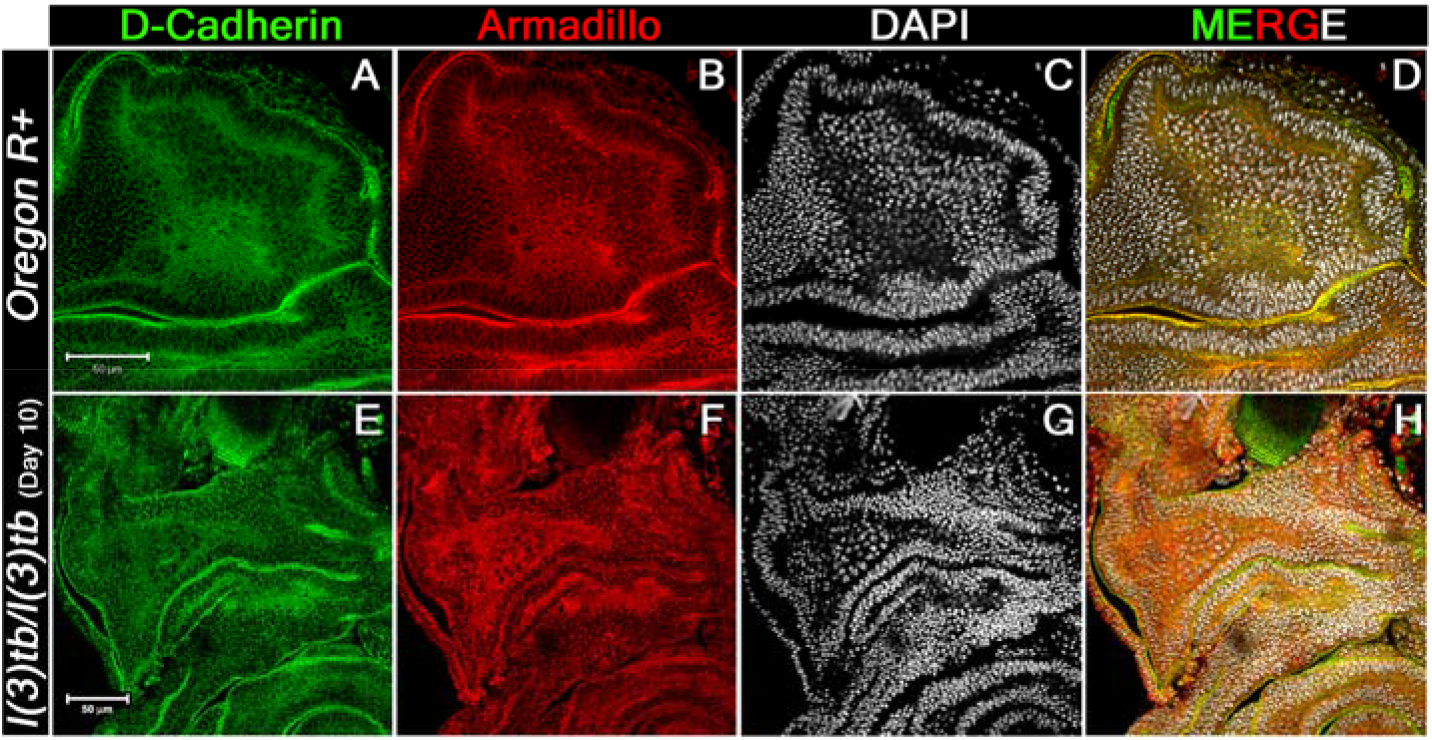
Magnified image of notum region in 3^rd^ instar larval wing imaginal discs immunolabeled to visualize the distribution pattern of DE-cadherin and □□Catenin. Distinct pattern of DE-cadherin (A) and Armadillo (B) immunostaining is visible in the wild type wing imaginal discs (notum region), which is absent in *l(3)tb* mutant wing imaginal disc (E & F respectively). The tissues were counterstained with DAPI (white). Scale bar is indicative of 50 μm.

**FIGURE S4.**
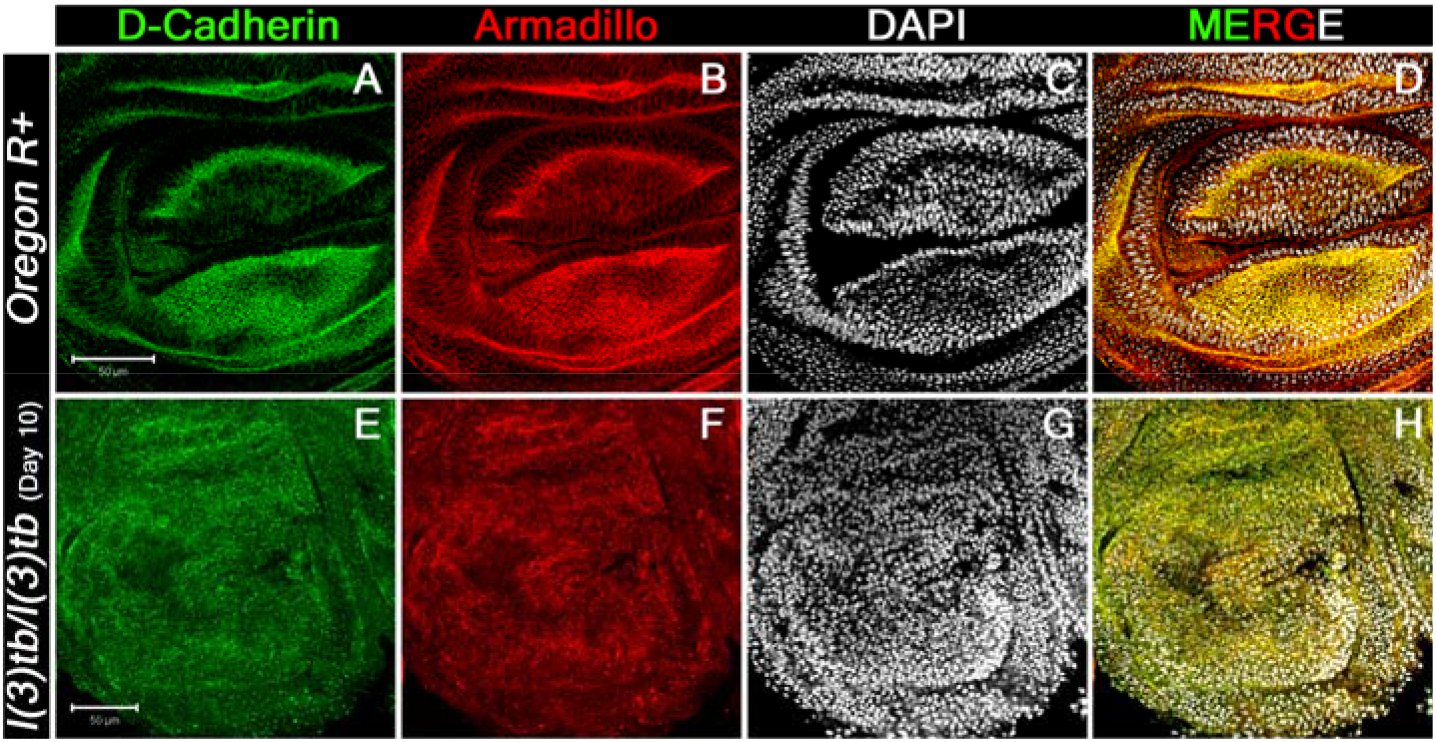
Magnified image of pouch region in 3^rd^ instar larval wing imaginal discs immunolabeled to visualize the distribution pattern of D-cadherin and □□Catenin. Cadherin staining was done with anti-DCadherin antibody (green) and □-Catenin staining was done with anti-Armadillo antibody (red) followed by counterstaining with DAPI (white). Scale bar indicates 50 μm.

**FIGURE S5.**
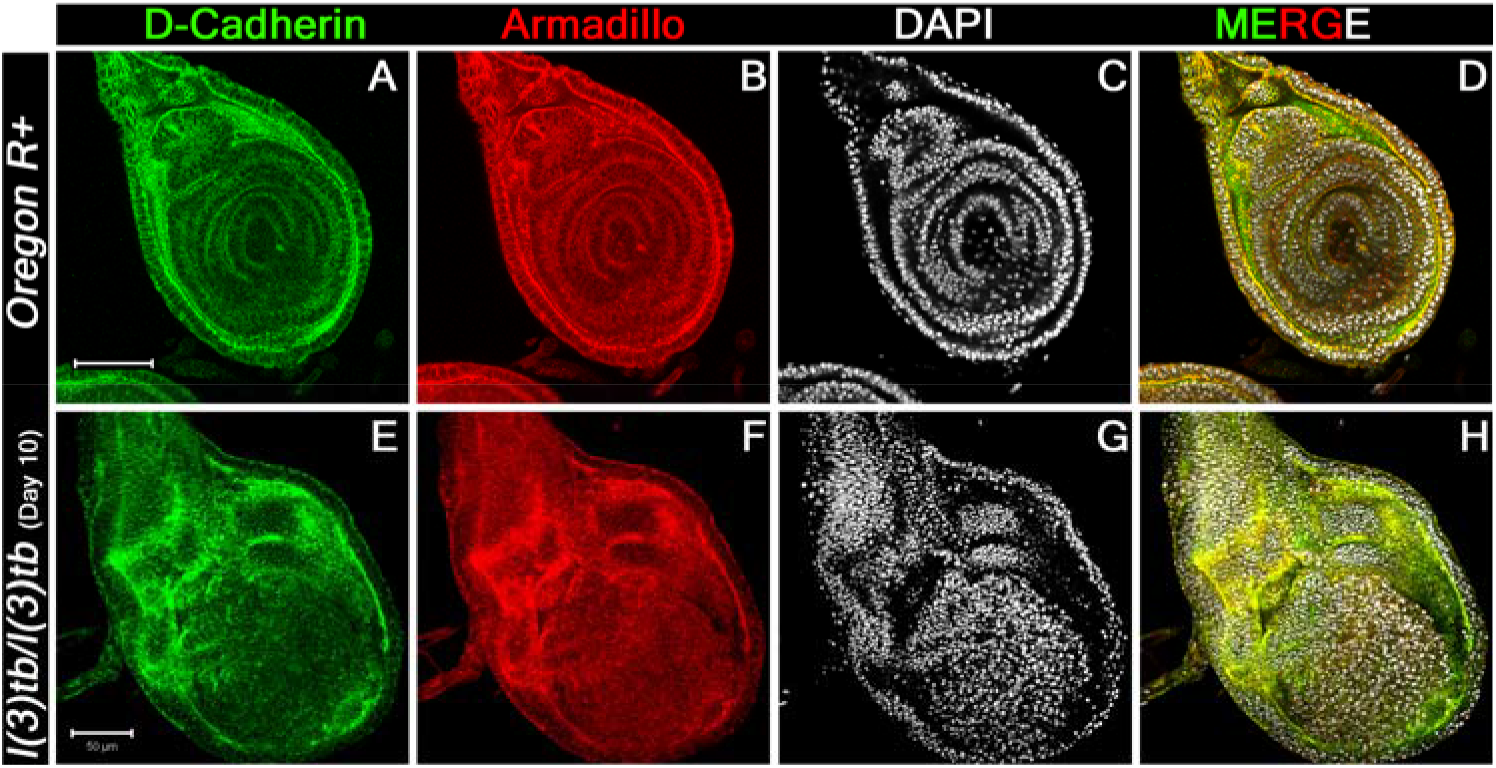
Confocal images of 3^rd^ instar larval leg imaginal discs immunolabeled to visualize the distribution pattern of DE-cadherin-catenin (green) complex proteins. Armadillo (□-Catenin, red) is a binding partner of trans-membranous protein DE-cadherin (green) having roles in cell adhesion and regulate tissue organization and morphogenesis. Scale bar indicates 50 μm

**FIGURE S6.**
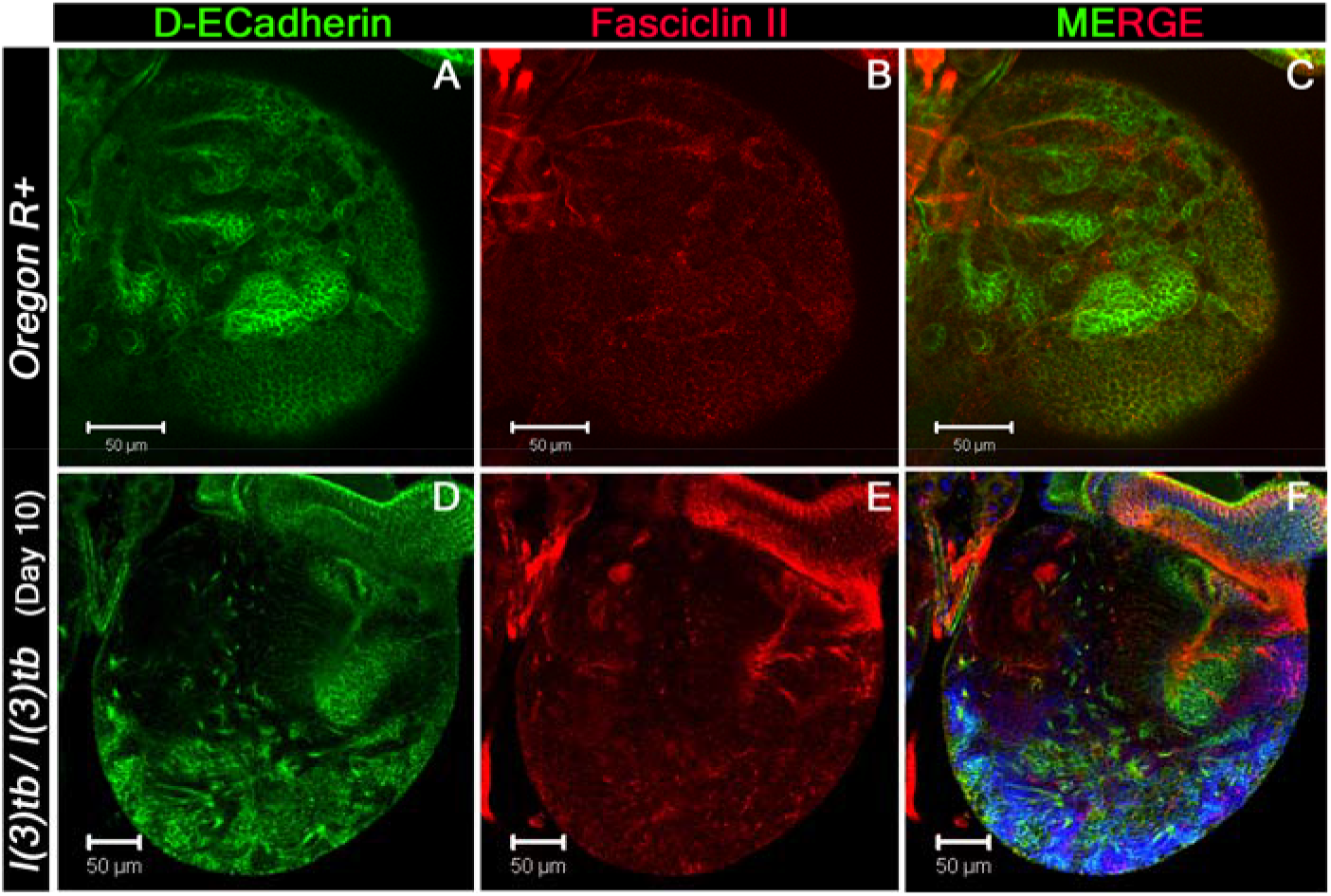
Confocal image of 3^rd^ instar larval brain lobes extending the distribution pattern of DE-cadherin (Green) and Fasciclin II (red). The pattern shown by DE cadherin, which labels the surface glia and neuropile glia, is altogether altered in the homozygous *l(3)tb* mutant (D) as compared to wild type (A) which shows strong staining in neuropile glia. Fas II labels the pioneering axonal fascicles in wild type (B) as shown by the marked mushroom body, MB, which is absent in the homozygous *l(3)tb* mutant (E). Fas II Images in C and E demonstrate the merged images of wild type and mutant respectively. Scale bar is 50 μm.

**FIGURE S7.**
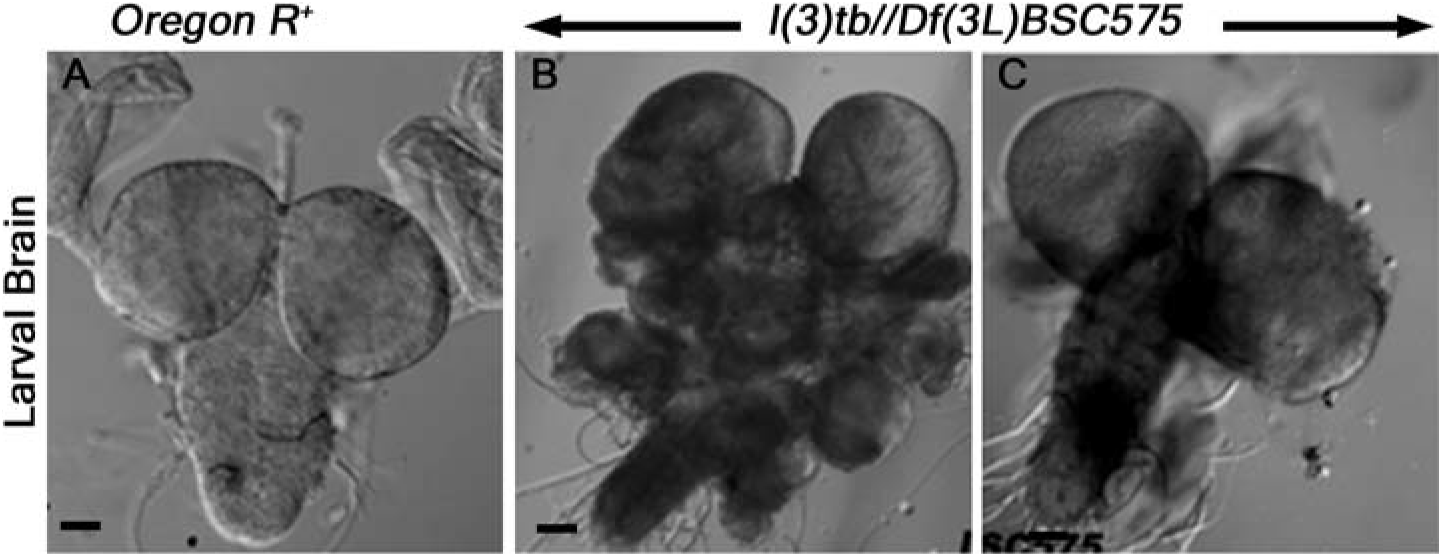
Tumorous larval brain phenotype in trans-heterozygous *l(3)tb* with *Df(3L)BSC575*. The deletion line, *Df(3L)BSC575*, do not show complementation with *l(3)tb* and the larvae showed tumorous brain in trans-heterozygous condition (B, C) when compared to wild type (A).

**FIGURE S8.**
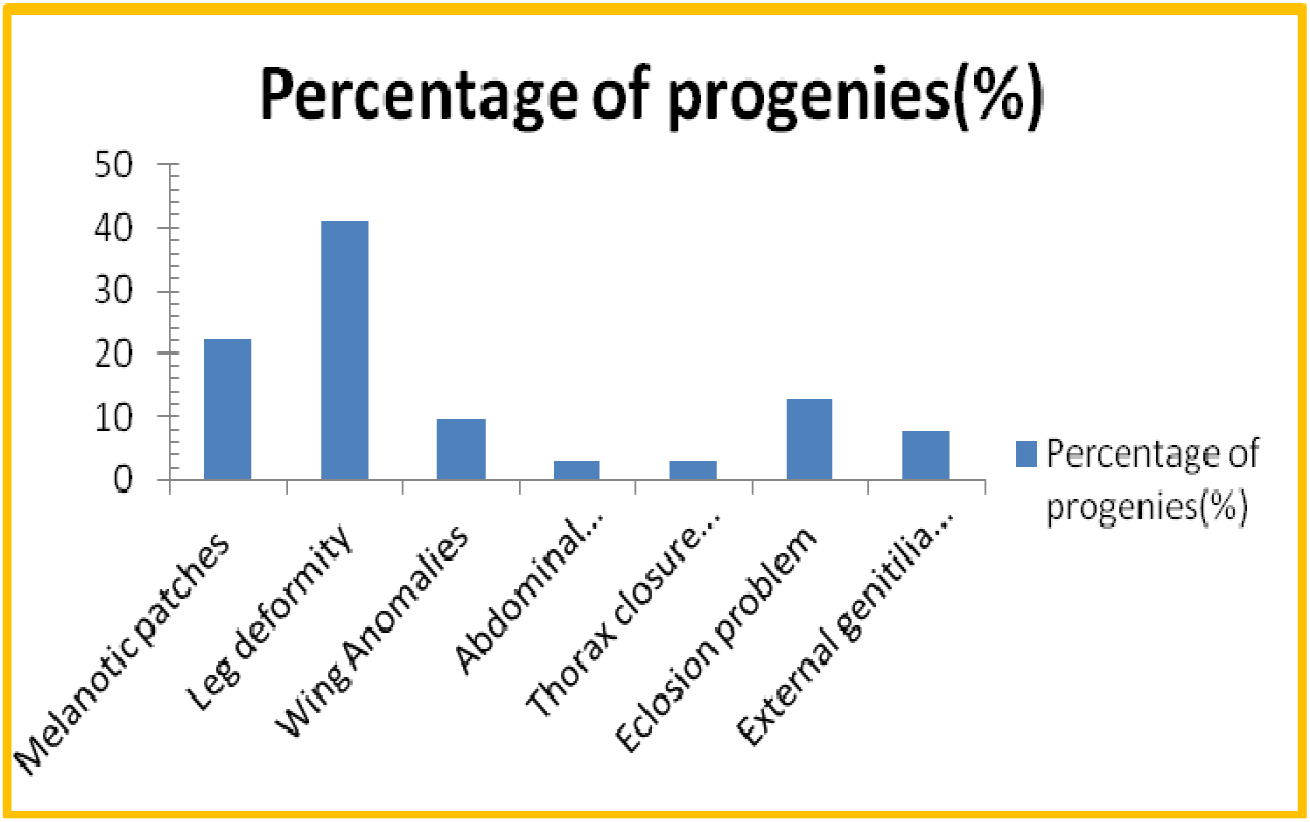
Morphological defects exhibited by escapees of adult fly trans-heterozygous for *P{GT1}DCP2^BG01766^/l(3)tb*. The phenotype includes melanotic patches (22.2%) on the cuticular exoskeleton, abnormalities in leg (41.3%), wing (10%), abdomen (3.2%) and thorax (3.2%). Many of the trans-heterozygous progeny was observed to have eclosion problem (12.7%) and males have abnormal genitalia (9.7%).

**FIGURE S9.**
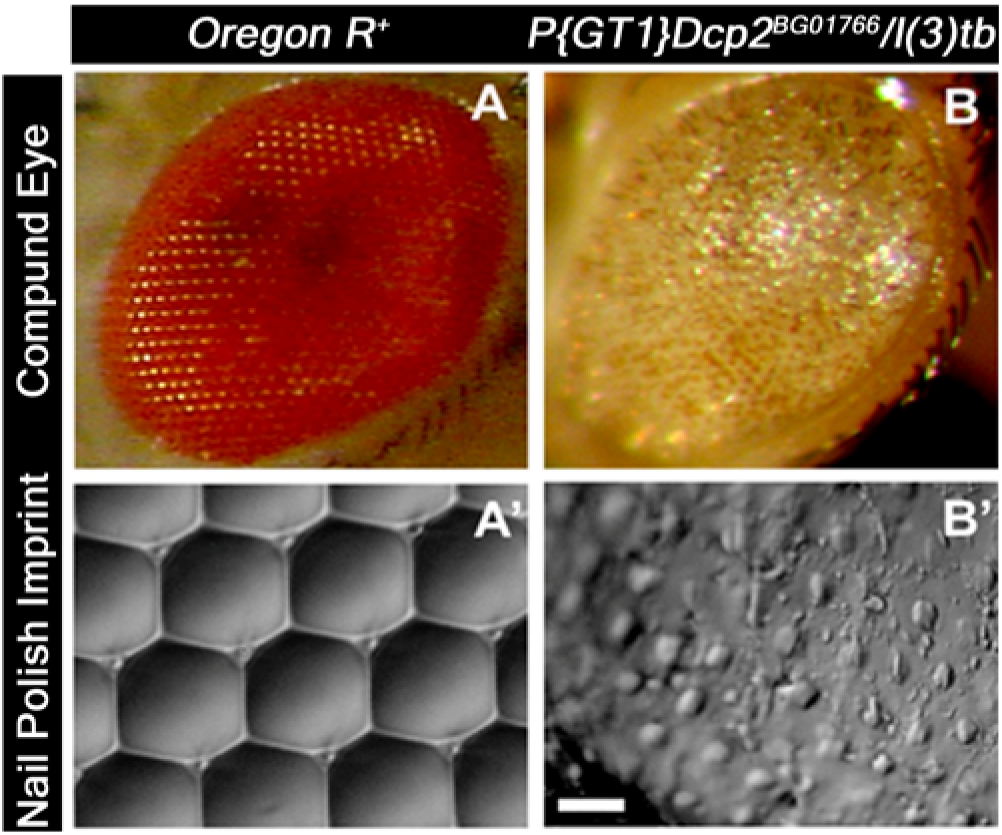
Pronouncement of severe defects in compound eyes of the escapees having heterozygous genetic background of the mutant *l(3)tb* with lethal P-insertion allele *DCP2^BG01766^*. Images in A and B showing the compound eye of wild type and tans-heterozygote respectively while A’ and B’ are their respective nail-polish imprint of the compound eye, viewed with the help of DIC or Nomarski microscope. The exact geometrical arrangement of ommatidia in a hexagonal pattern having each ommatidium surrounded by bristle was completely disrupted in the trans-heterozygote exhibiting the complete loss of arrangement in the ommatidial pattern. This represents the severe loss of polarity as it cues a complete disassembly of compound eye as whole. Bar represents 20μm.

**FIGURE S10.**
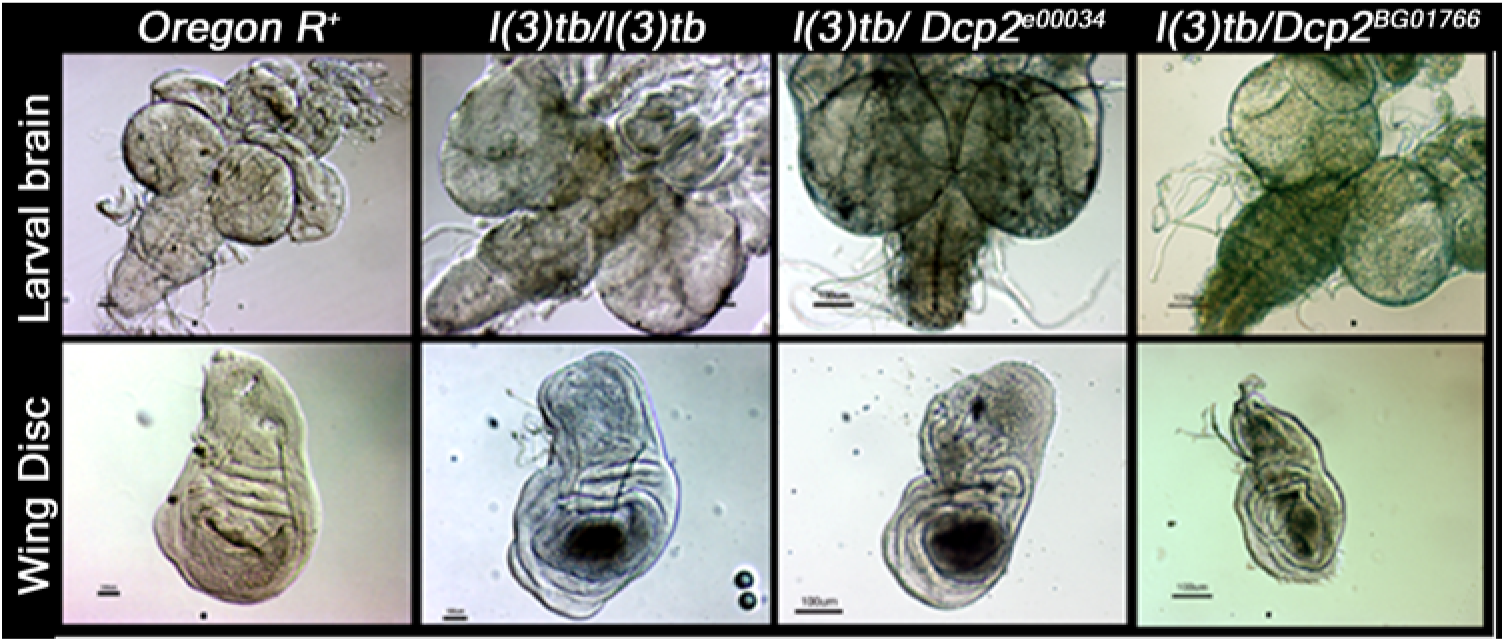
Tumorous phenotype observed in larval brain and wing imaginal discs in transheterozygotes *l(3)tb /PBac{RB}DCP2^e00034^* and *l(3)tb /P{GT1}DCP2^BG01766^* as homozygous *l(3)tb* Scale bar is 100μm.

**FIGURE 11.**
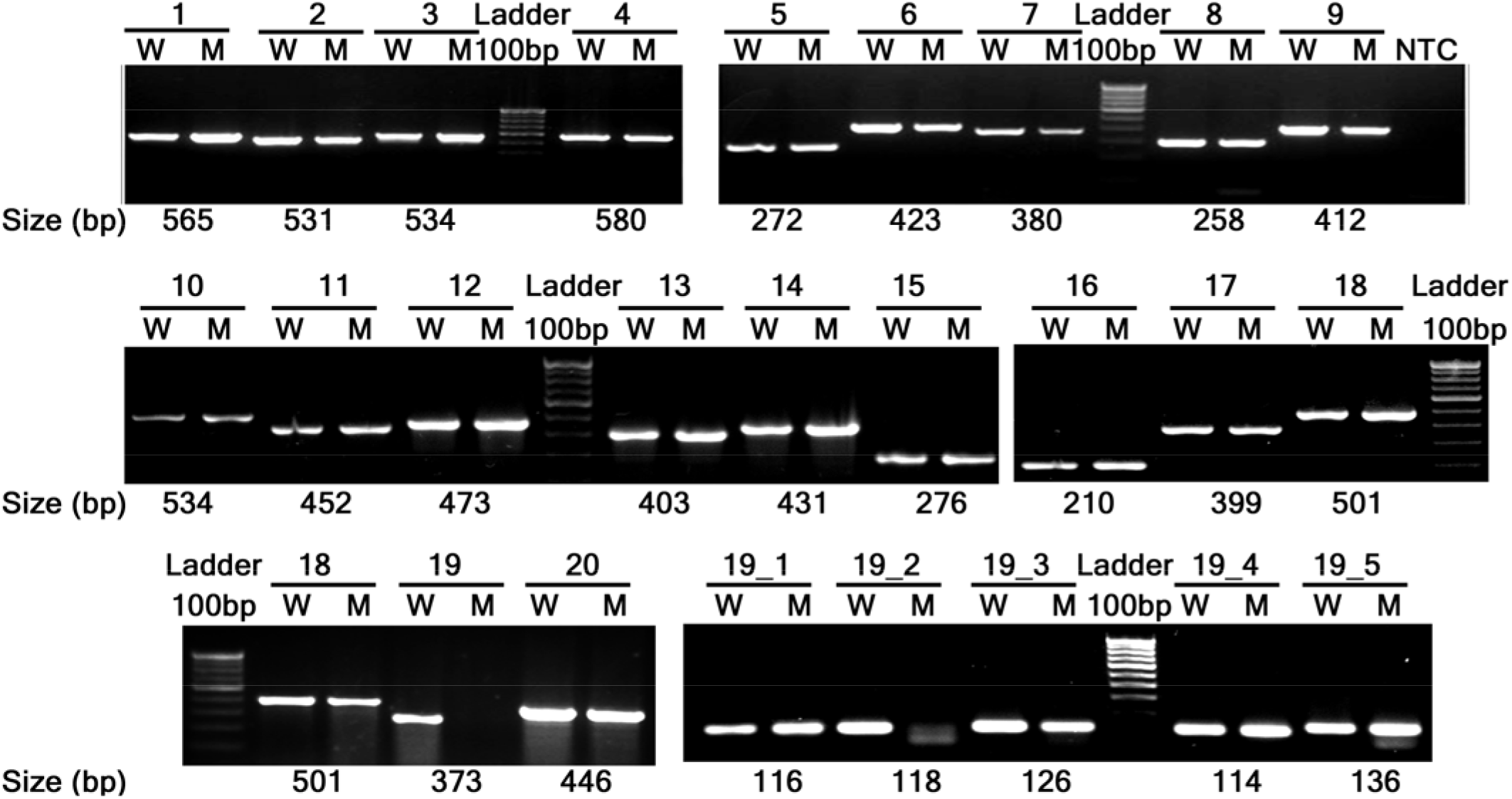
Amplification of DCP2 gene using overlapping primers. All primers amplify same size of amplicon with DNA from wild type and homozygous l(3)tb mutant, except DCP2_P19 (3L: 15819379.15819751) and DCP2_P19_2 (3L:15819452.15819569). This implies the probable mutation in the region.

**TABLE S1.** Various deletion lines used for complementation with the mutation in *l(3)tb*. All the below deletion lines complemented with the mutation in *l(3)tb* except *Df(3L)BSC845, Df(3L)BSC774* and *Df(3L)BSC575* which resulted in non-complementation to the mutation in *l(3)tb*. [* = marked *Df(3L)RM95*, molecularly characterized deletion generated in the laboratory and also exhibiting noncomplementation].

| S. No. | Deletion Lines | Estimated Cytology | Genomic Sequence Co-ordinates of Break points (Proximal; Distal) | Deleted Region (in base) | Status of Complementation |
| --- | --- | --- | --- | --- | --- |
| 1. | *Df(3L)ED4079* | 61A5;61B1 | 3L:40319; 131780 | 715336 | Yes |
| 2. | *Df(3L)ED201* | 61B1;61C1 | 3L:123924; 347941 | 347941 | Yes |
| 3. | *Df(3L)ED4177* | 61C1;61E2 | 3L:319846; 1035182 | 1035182 | Yes |
| 4. | *Df(3L)ED4287* | 62B4;62E5 | 3L:1795442; 2551761 | 756319 | Yes |
| 5. | *Df(3L)ED4288* | 63A6;63B7 | 3L:3070827; 3149091 | 78264 | Yes |
| 6. | *Df(3L)ED208* | 63C1;63F5 | 3L:3249148; 3893148 | 644000 | Yes |
| 7. | *Df(3L)ED4341* | 63F6;64B9 | 3L:3905091; 4542236 | 637145 | Yes |
| 8. | *Df(3L)ED210* | 64B9;64C13 | 3L:4544234; 5348442 | 804208 | Yes |
| 9. | *Df(3L)ED211* | 65A9;65B4 | 3L:6211235; 6545859 | 334624 | Yes |
| 10. | *Df(3L)ED4408* | 66A22;66C5 | 3L:7972207; 8292674 | 320467 | Yes |
| 11. | *Df(3L)ED4421* | 66D12;67B3 | 3L:8738426; 9377175 | 638749 | Yes |
| 12. | *Df(3L)ED4457* | 67E2;68A7 | 3L:10357051; 11118909 | 761858 | Yes |
| 13. | *Df(3L)ED4470* | 68A6;68E1 | 3L:11090089; 11826330 | 736241 | Yes |
| 14. | *Df(3L)ED4475* | 68C13;69B4 | 3L:11580140; 12401701 | 821561 | Yes |
| 15. | *Df(3L)ED4483* | 69A5;69D3 | 3L:12270320; 12686314 | 415994 | Yes |
| 16. | *Df(3L)ED4486* | 69C4;69F6 | 3L:12507519; 13025585 | 518066 | Yes |
| 17. | *Df(3L)ED4502* | 70A3;70C10 | 3L:13220865; 13986651 | 765786 | Yes |
| 18. | *Df(3L)ED4543* | 70C6;70F4 | 3L:13928325; 14751140 | 822815 | Yes |
| 19. | *Df(3L)ED217* | 70F4;71E1 | 3L:14751170; 15582196 | 831026 | Yes |
| 20. | *Df(3L)BSC833* | 71D4;71F1 | 3L:15507245; 15671057 | 163812 | Yes |
| 21. | *Df(3L)RM95\** | 71E1;72B2 | 3L:15481449; 15904040 | 422591 | NO |
| 22. | *Df(3L)BSC845* | 71D3;72A1 | 3L:15504128; 15819023 | 314895 | NO |
| 23. | *Df(3L)BSC575* | 71F1;72C1 | 3L:15671057; 15973064 | 302007 | NO |
| 24. | *Df(3L)BSC774* | 71F1;72D10 | 3L:15693003; 16233380 | 540377 | NO |
| 25. | *Df(3L)Exel6127* | 72D1;72D9 | 3L:16040040; 16122754 | 82714 | Yes |
| 26. | *Df(3L)ED223* | 73A1;73D5 | 3L:16444925; 16883977 | 439052 | Yes |
| 27. | *Df(3L)ED4674* | 73B5;73E5 | 3L:16654384; 17042518 | 388134 | Yes |
| 28. | *Df(3L)ED4685* | 73D5;74E2 | 3L:16884176; 17605270 | 721094 | Yes |
| 29. | *Df(3L)ED4710* | 74D1;75B11 | 3L:17480563; 18132399 | 651836 | Yes |
| 30. | *Df(3L)ED224* | 75B1;75C6 | 3L:17962303; 18391619 | 429316 | Yes |
| 31. | *Df(3L)ED225* | 75C1;75D4 | 3L:18179245; 18614437 | 435192 | Yes |
| 32. | *Df(3L)ED4782* | 75F2;76A1 | 3L:18988994; 19163802 | 174808 | Yes |
| 33. | *Df(3L)ED4786* | 75F7;76A5 | 3L:19094051; 19288762 | 194711 | Yes |
| 34. | *Df(3L)ED229* | 76A1;76E1 | 3L:19163806; 19995811 | 832005 | Yes |
| 35. | *Df(3L)ED4858* | 76D3;77C1 | 3L:19888473; 20394920 | 506447 | Yes |
| 36. | *Df(3L)ED4978* | 78D5;79A2 | 3L:21526907; 21873785 | 346878 | Yes |
| 37. | *Df(3L)ED230* | 79C2;80A4 | 3L:22127751; 22827471 | 699720 | Yes |
| 38. | *Df(3L)ED5017* | 80A4;80C2 | 3L:22828597; 22991401 | 162804 | Yes |
| 39. | *Df(3R)ED5100* | 82A1;82E8 | 3R:22995; 912807 | 889812 | Yes |
| 40. | *Df(3R)ED5138* | 82D5;82F8 | 3R:606794; 1090605 | 483811 | Yes |
| 41. | *Df(3R)ED5142* | 82B3;82F8 | 3R:279018; 1090605 | 811587 | Yes |
| 42. | *Df(3R)ED5147* | 82E8;83A1 | 3R:912842; 1193526 | 280684 | Yes |
| 43. | *Df(3R)ED5156* | 82F8;83A4 | 3R:1090655; 1284574 | 193919 | Yes |
| 44. | *Df(3R)ED10257* | 83A7;83B4 | 3R:1344078; 1426000 | 81922 | Yes |
| 45. | *Df(3R)ED5177* | 83B4;83B6 | 3R:1426351; 1449817 | 23466 | Yes |
| 46. | *Df(3R)ED5197* | 83B7;83D2 | 3R:1474504; 1833866 | 359362 | Yes |
| 47. | *Df(3R)ED7665* | 84B4;84E11 | 3R:2916249; 3919805 | 1003556 | Yes |
| 48. | *Df(3R)ED5230* | 84E6;85A5 | 3R:3803496; 4478856 | 675360 | Yes |
| 49. | *Df(3R)ED5296* | 84F6;85C3 | 3R:4076143; 4882413 | 806270 | Yes |
| 50. | *Df(3L)ED5330* | 85A5;85D1 | 3R:4495308; 5055517 | 560209 | Yes |
| 51. | *Df(3R)ED5339* | 85D1;85D11 | 3R:5052798; 5178097 | 125299 | Yes |
| 52. | *Df(3R)ED5429* | 85D21;85F8 | 3R:5336031; 5874333 | 538302 | Yes |
| 53. | *Df(3L)ED5454* | 85E5;85F12 | 3R:5552399; 5937180 | 384781 | Yes |
| 54. | *Df(3R)ED5474* | 85F11;86B1 | 3R:5935134; 6176446 | 241312 | Yes |
| 55. | *Df(3L)ED5495* | 85F16;86C7 | 3R:5996223; 6712482 | 716259 | Yes |
| 56. | *Df(3R)ED5559* | 86E11;87B11 | 3R:7394904; 8269738 | 874834 | Yes |
| 57. | *Df(3L)ED5577* | 86F9;87B13 | 3R:7654463; 8303300 | 648837 | Yes |
| 58. | *Df(3L)ED5591* | 87B7;87C7 | 3R:8176253; 8545732 | 369479 | Yes |
| 59. | *Df(3R)ED5610* | 87B11;87D7 | 3R:8269738; 8821397 | 551659 | Yes |
| 60. | *Df(3L)ED5623* | 87E3;88A4 | 3R:9085471; 9809634 | 724163 | Yes |
| 61. | *Df(3R)ED5642* | 87F10;88C2 | 3R:9509544; 10307496 | 797952 | Yes |
| 62. | *Df(3R)ED5644* | 88A4;88C9 | 3R:9843625; 10451431 | 607806 | Yes |
| 63. | *Df(3R)ED10639* | 89B7;89D2 | 3R:12038635; 12306942 | 268307 | Yes |
| 64. | *Df(3R)ED10635* | 89B13;89D2 | 3R:12174977; 12306942 | 131965 | Yes |
| 65. | *Df(3R)ED10642* | 89C7;89D5 | 3R:12279479; 12450993 | 171514 | Yes |
| 66. | *Df(3L)ED5780* | 89E11;90C1 | 3R:12882199; 13507523 | 625324 | Yes |
| 67. | *Df(3R)ED5785* | 90C2;90D1 | 3R:13543832; 13769792 | 225960 | Yes |
| 68. | *Df(3L)ED5797* | 90C2;90F10 | 3R:13543832; 14068391 | 524560 | Yes |
| 69. | *Df(3R)ED5815* | 90F4;91B8 | 3R:13993596; 14484708 | 491112 | Yes |
| 70. | *Df(3R)ED2* | 91A5;91F1 | 3R:14224953; 14922493 | 697540 | Yes |
| 71. | *Df(3R)ED5911* | 91C5;91F8 | 3R:14568649; 14991505 | 422856 | Yes |
| 72. | *Df(3R)ED6025* | 92A11;92E2 | 3R:15468450; 16135241 | 666791 | Yes |
| 73. | *Df(3R)ED10820* | 93A4;93B12 | 3R:16774462; 16937182 | 162720 | Yes |
| 74. | *Df(3R)ED10845* | 93B9;93D4 | 3R:16890893; 17122221 | 231328 | Yes |
| 75. | *Df(3R)ED6052* | 93D4;93D8 | 3R:17122205; 17191074 | 68869 | Yes |
| 76. | *Df(3R)ED6076* | 93E10;94A1 | 3R:17459227; 17868550 | 409323 | Yes |
| 77. | *Df(3L)ED6085* | 93F14;94B5 | 3R:17706717; 18413461 | 706744 | Yes |
| 78. | *Df(3R)ED6096* | 94B5;94E7 | 3R:18413403; 19047691 | 634288 | Yes |
| **79.** | *Df(3R)ED6103* | 94D3;94E9 | 3R:18724275; 19084137 | 359862 | Yes |
| **80.** | *Df(3R)ED6187* | 95D10;96A7 | 3R:19877370; 20369665 | 492295 | Yes |
| **81.** | *Df(3R)ED6220* | 96A7;96C3 | 3R:20369520; 21009495 | 639975 | Yes |
| **82.** | *Df(3R)ED6235* | 97B9;97D12 | 3R:22360956; 22806229 | 445273 | Yes |
| **83.** | *Df(3R)ED6255* | 97D2;97F1 | 3R:22624758; 23107623 | 482865 | Yes |
| **84.** | *Df(3R)ED6265* | 97E2;98A7 | 3R:22937981; 23405492 | 467511 | Yes |
| **85.** | *Df(3R)ED6310* | 98F12;99B2 | 3R:24964617; 25337875 | 373258 | Yes |
| **86.** | *Df(3R)ED6316* | 99A5;99C1 | 3R:25081045; 25608389 | 527344 | Yes |

**TABLE S2.**
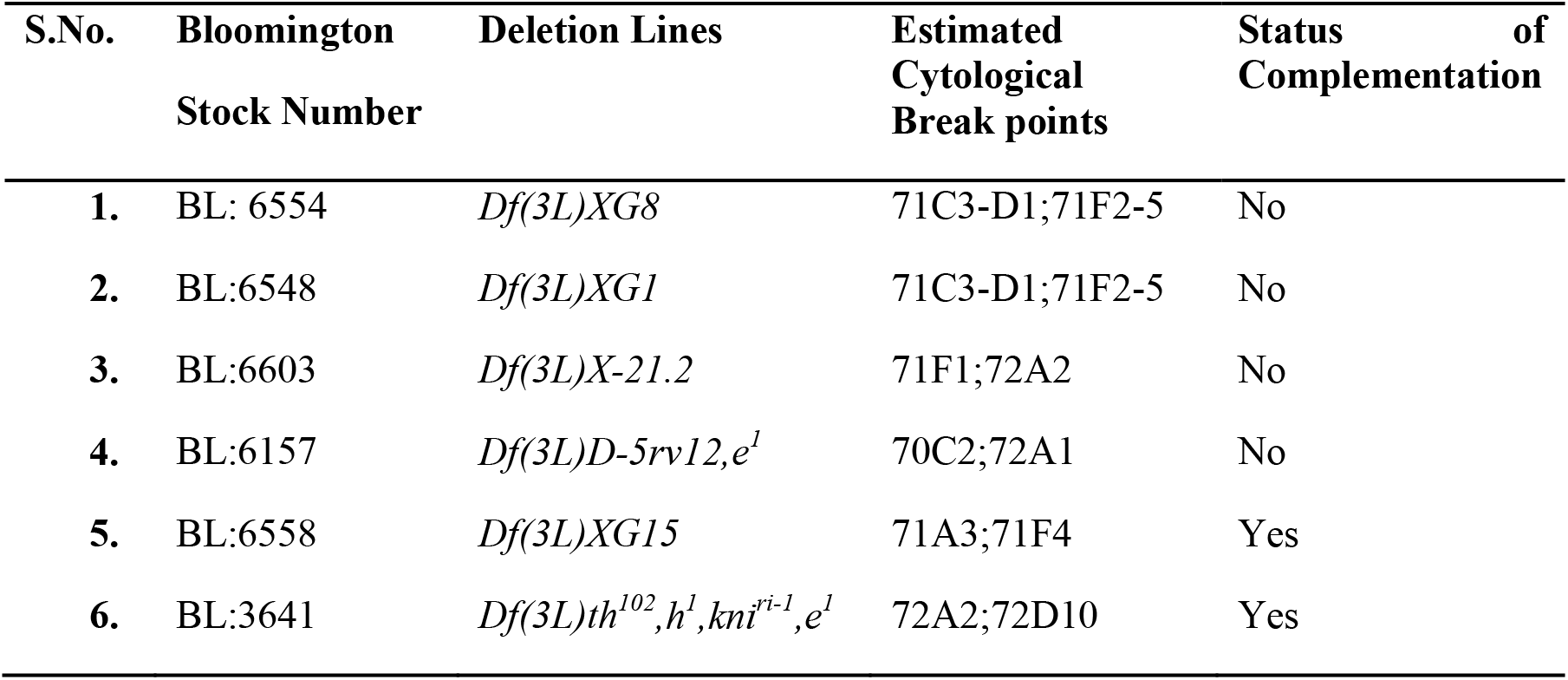
Complementation status of *l(3)tb* with cytologically mapped deletion lines

**TABLE S3.**
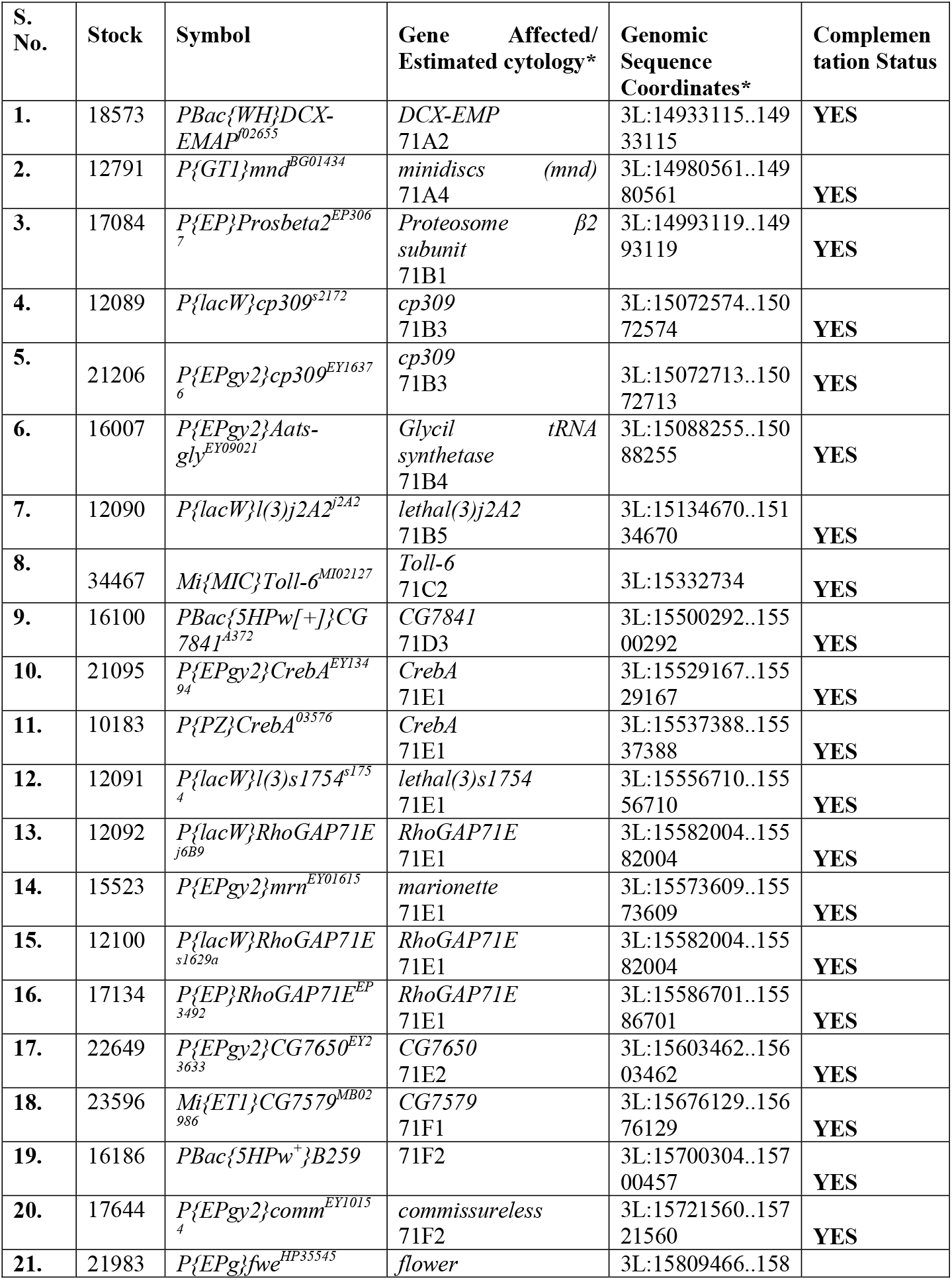
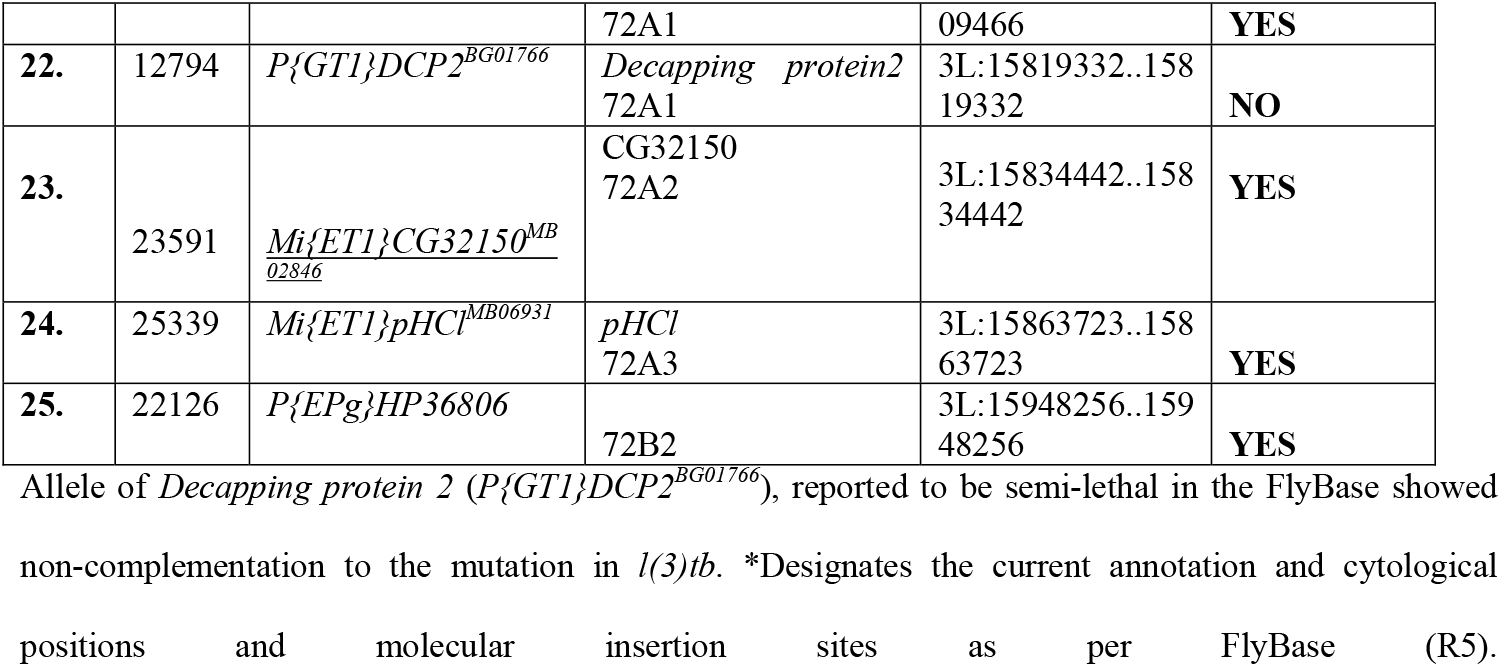
Complementation analysis of *l(3)tb* with lethal transposon insertion lines

**TABLE S4.**
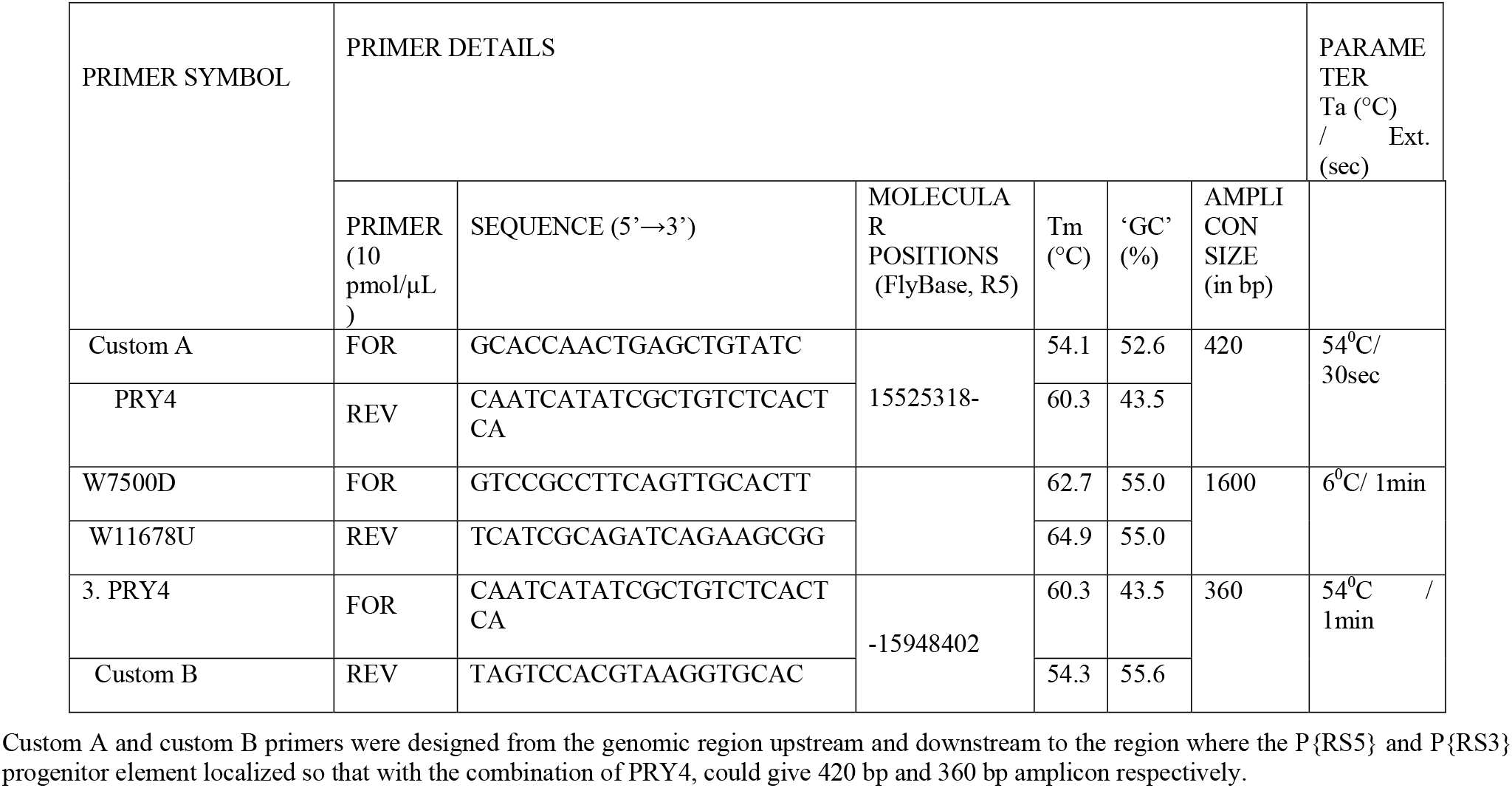
Primers used for characterizing deletion in *Df(3L)RM95*.

**TABLE S5.**
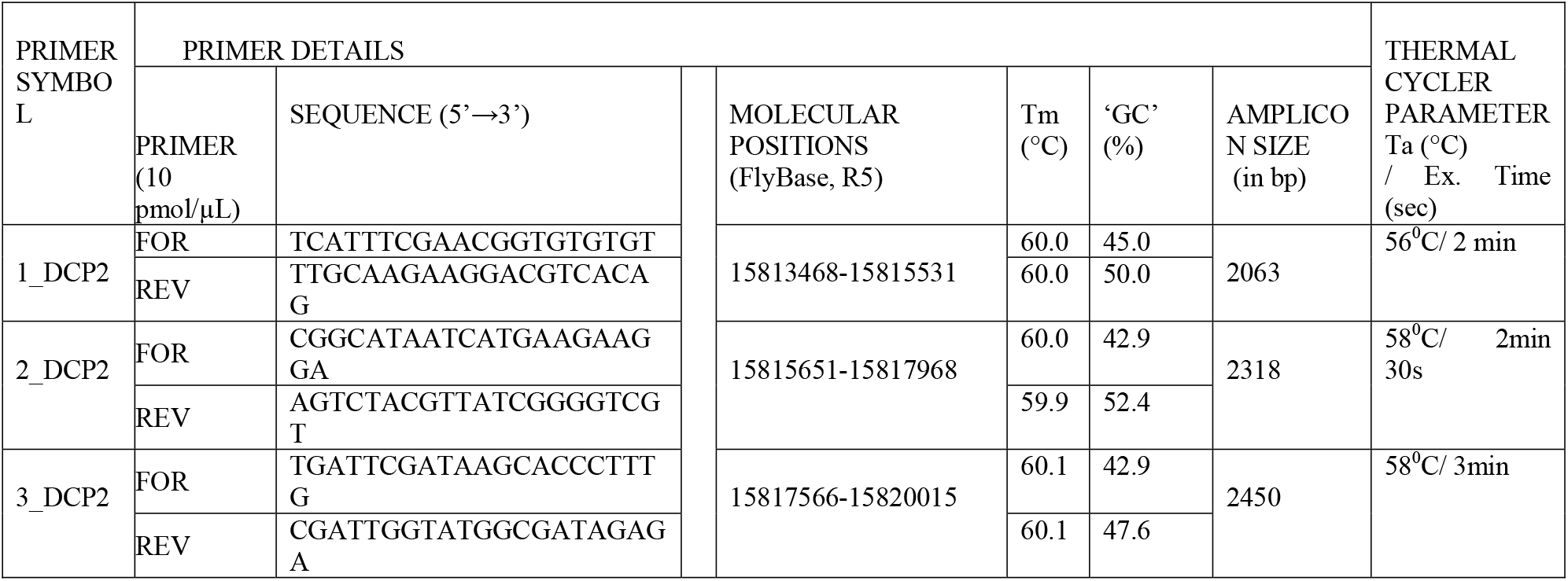
First set of primers used for amplification of genomic region of *Decapping protein 2* in the mutant *l(3)tb*.

**TABLE S6.**
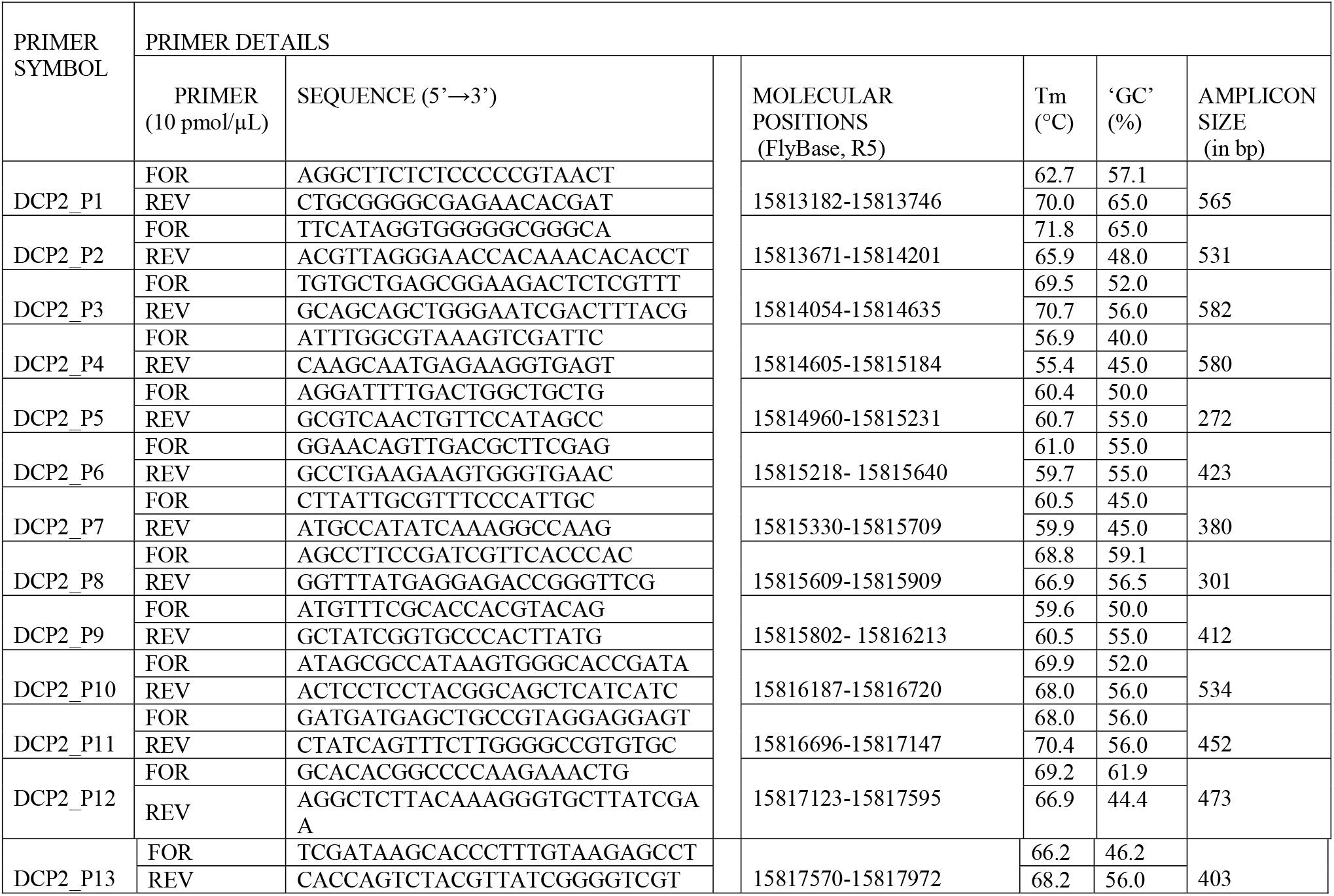
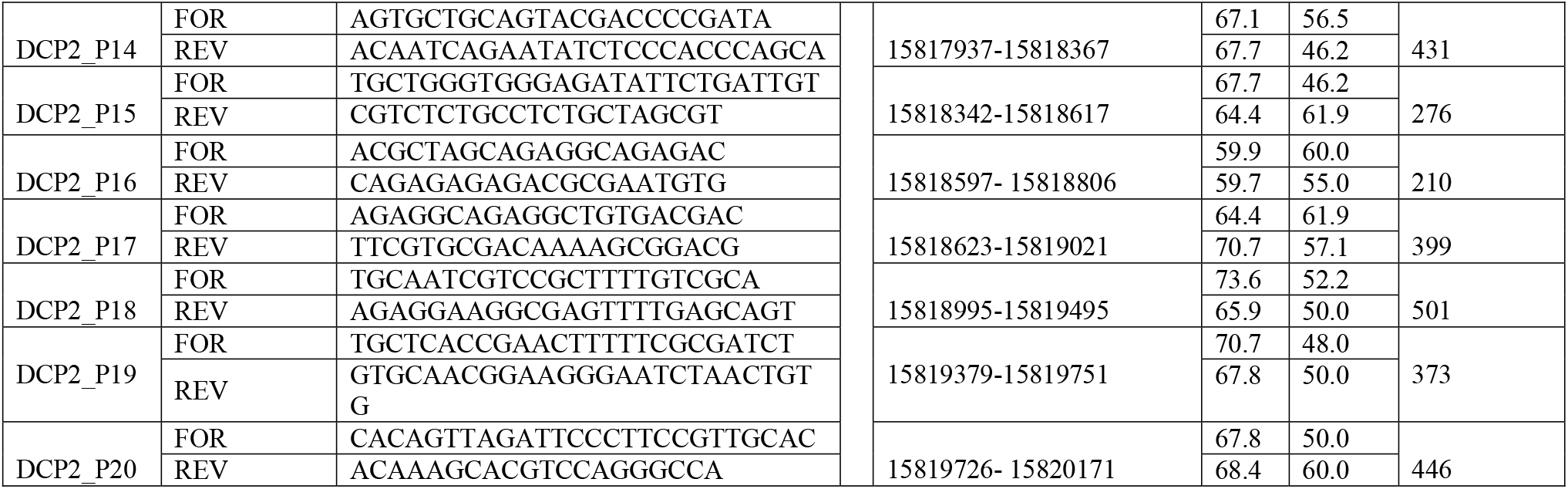
Overlapping set of primers for *DCP2* gene and thermal cycler conditions of annealing temperature and extension time for each primer pair to amplify the genomic region of *DCP2* gene in the homozygous *l(3)tb* mutant.

**TABLE S7.**
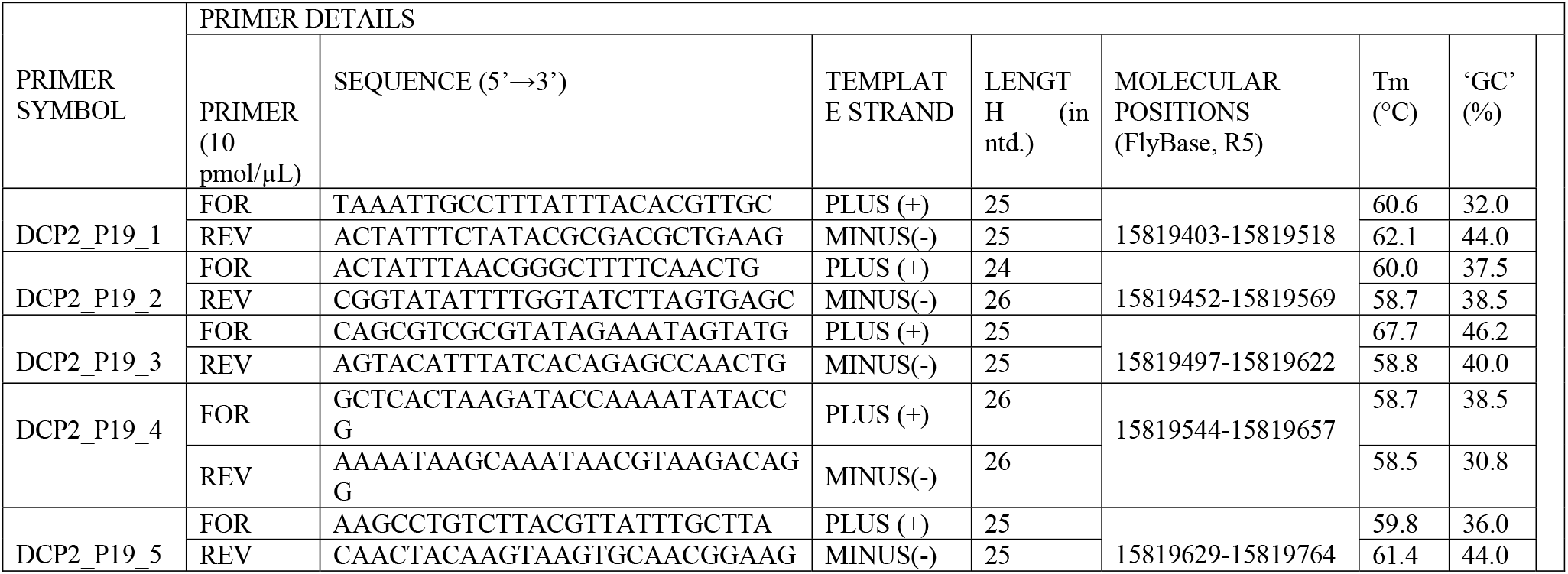
Overlapping set of primers to amplify the genomic region in *DCP2* gene for the region covered by the DCP2_P19 set of primers in the homozygous *l(3)tb* mutant.

**TABLE S8.**
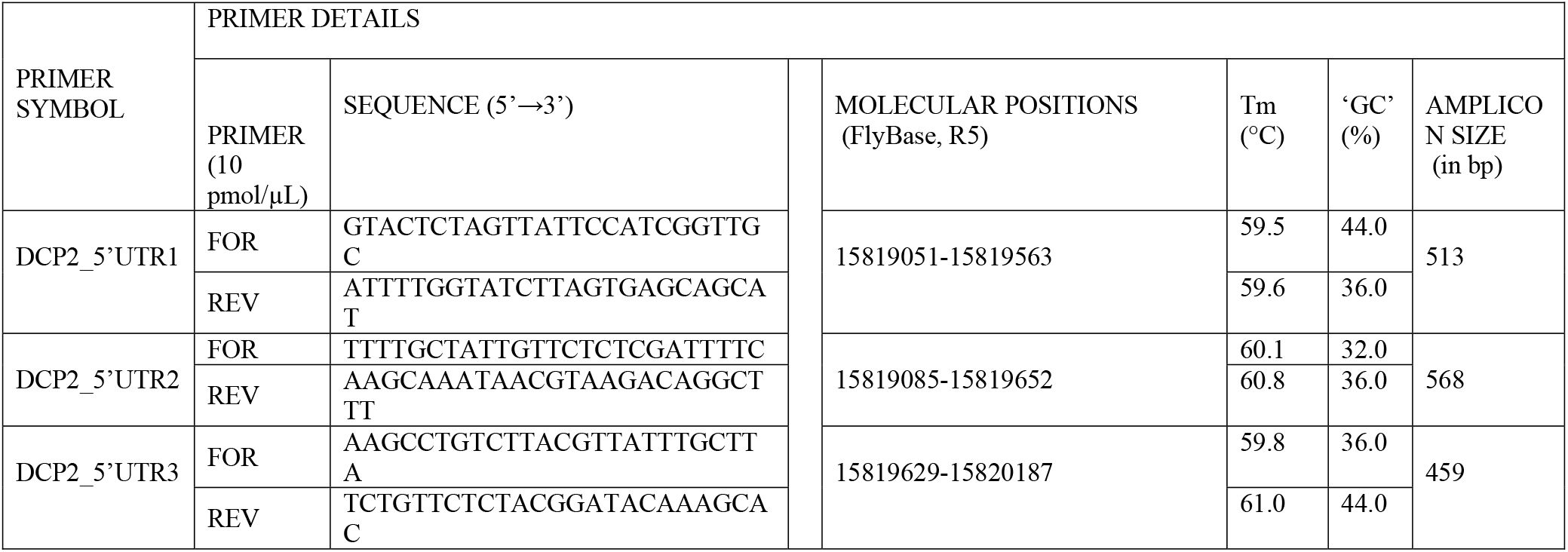
Overlapping set of primers to amplify the complete 5’UTR of genomic region in *DCP2* gene in the homozygous *l(3)tb* mutant. The table also documents the thermal cycler conditions of annealing temperature and extension time for each primer pair. Genomic region amplified by primer pair is also mentioned.

**TABLE S9.** Three promoters for *Drosophila* gene *DCP2* are shown to provide transcriptional regulation for six transcripts.

| Locus | Promoter ID/ | Strand | Length | Transcript/TSS |
| --- | --- | --- | --- | --- |
| **Decapping Protein 2, DCP2 (GeneID:39722)**<br>**DmelCG6169**<br>**GXL_196353**<br>**NCBI Build 5.41**<br>**Chromosome 3L**<br>**Contig NT_037436** | GXP_235577<br>15817463-15818063<br>1 relevant transcript | Minus | 601 bp | GXT_1658903 (5 Exons)<br>NM_206378 (DCP2-RC)***<br>TSS=501 |
|  | GXP_622284<br>15817204-15817804<br>2 relevant transcript | Minus | 601 bp | GXT_22014755 (5 Exons)<br>NM_001014585 (DCP2-RD)<br>TSS=501<br>GXT_25225584 (6 Exons)<br>FBtr0100528 (DCP2-RD)<br>TSS=501 |
|  | GXP_622285<br>15819404-15820020<br>3 relevant transcript | Minus | 617 bp | GXT_22014753 (5 Exons)<br>NM_140548 (DCP2-RA)<br>TSS=502<br>GXT_22014754 (5 Exons)<br>NM_168634 (DCP2-RB)<br>TSS=517<br>GXT_25225583 (5 Exons)<br>FBtr0304975 (DCP2-RE)<br>TSS=501 |

\*\*\* The NM\_206378 transcript has been removed from the database. (www.genomatix.de )

